# Structural determinants of co-translational protein complex assembly

**DOI:** 10.1101/2024.01.20.576408

**Authors:** Saurav Mallik, Johannes Venezian, Arseniy Lobov, Meta Heidenreich, Hector Garcia-Seisdedos, Todd O. Yeates, Ayala Shiber, Emmanuel D. Levy

**Affiliations:** Department of Chemical and Structural Biology, Weizmann Institute of Science, Rehovot 7600001, Israel; Faculty of Biology, Technion-Israel Institute of Technology, Haifa 3200003, Israel; Department of Structural and Molecular Biology, Institute of Molecular Biology, Barcelona 08028, Spain; Department of Chemistry and Biochemistry, University of California, Los Angeles 90095, United States

## Abstract

The assembly of proteins into functional complexes is critical to life’s processes. While textbooks depict complex assembly as occurring between fully synthesized proteins, we know today that thousands of proteins in the human proteome assemble co-translationally during their synthesis. Why this process takes place, however, remains unknown. We show that co-translational assembly is governed by biophysical and structural characteristics of the protein complex, and involves mutually stabilized, intertwined subunits. Consequently, these subunits are also co-regulated across the central dogma, from transcription to protein degradation. Leveraging structural signatures with AlphaFold2-based predictions enables us to accurately predict co-translational assembly on a proteome-wide scale, which we validated by ribosome profiling, genetic perturbations, and smFISH experiments. Notably, the latter showed that co-translationally assembling subunits exhibit co-localized mRNAs. This work unveils a fundamental connection between protein structure and the translation process, highlighting the overarching impact of three-dimensional structure on gene expression, mRNA localization, and proteostasis.

**One Sentence Summary:** Protein complexes with topologically intertwined subunits require co-translational assembly and synchronized proteostasis of subunits, with implications in protein stability, mRNA localization, and evolution.

**Graphical Abstract:** 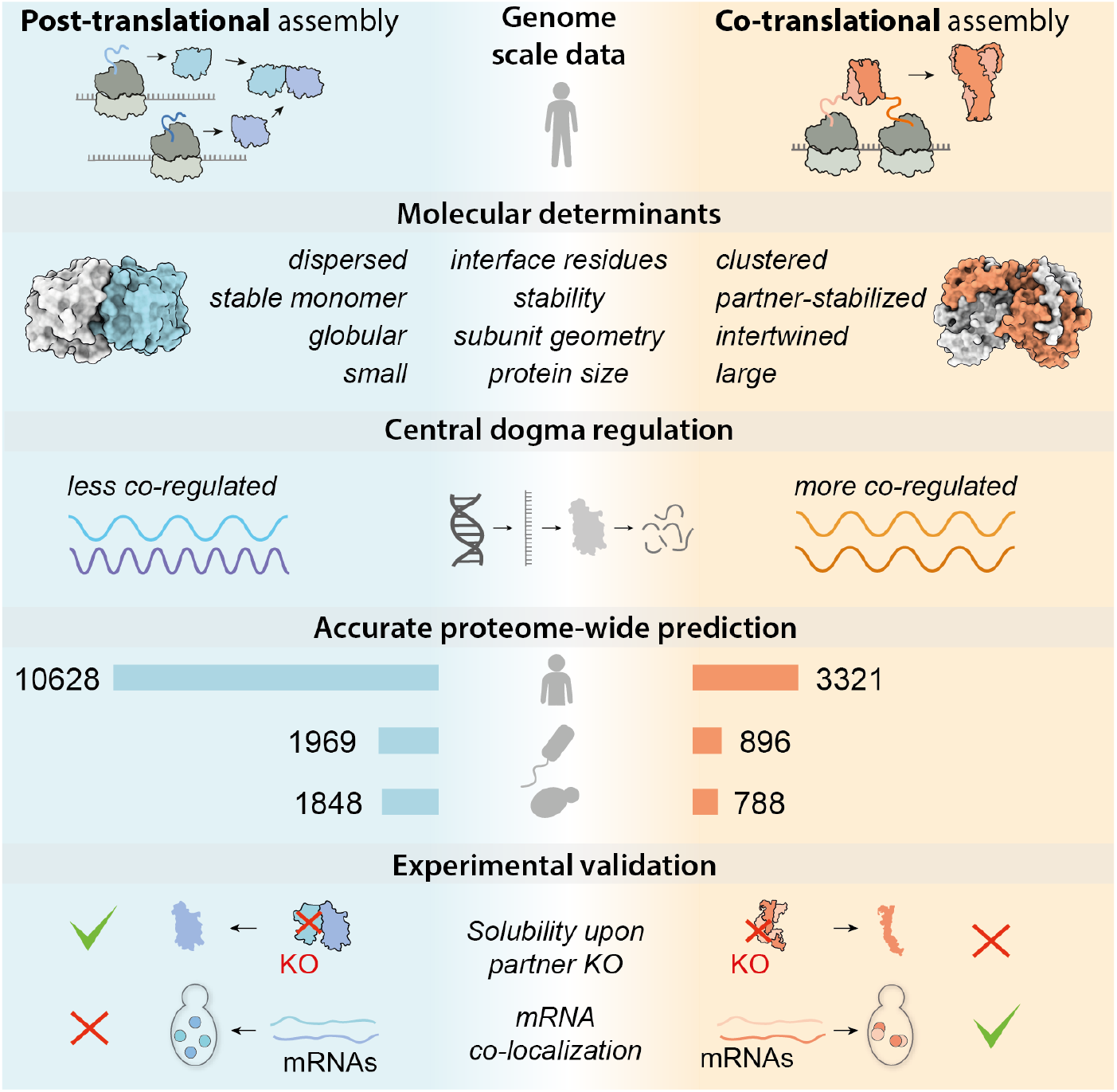

## INTRODUCTION

The self-organization of biological macromolecules is essential for executing cellular functions. Among proteins, self-organization ranges from folding into specific three-dimensional structures ^1–6^ to assembly into large multi-component complexes ^7–9^. Crucially, the ability to characterize these folding and assembly events in detail is intrinsically constrained by the suitability of proteins for biophysical analysis. Consequently, detailed biophysical studies of protein folding have predominantly concentrated on small, globular proteins due to their reversible folding properties ^6^. However, these proteins represent less than 10% of the structural diversity within even a bacterial proteome ^10^. Characterizing the remaining 90% poses a significant challenge, as these proteins are often more susceptible to misfolding or aggregation ^10–13^. This leaves the vast majority of the cellular machinery’s folding and assembly mechanisms largely unexplored.

The analysis of protein complexes’ assembly faces similar limitations. Indeed, the reconstitution of complexes can be notoriously challenging, necessitating specialized technologies for spatiotemporal control of gene expression ^11,12^. As a result, protein complex isolation often involves the purification of endogenously tagged copies and structural characterization, *e*.*g*., by cryo-electron microscopy (Cryo-EM) (16, 17). While Cryo-EM can provide insights into the assembly of complexes through snapshots acquired along their assembly pathway (12) or across different functional states (13, 14), such approaches are focused on specific systems so that molecular determinants of folding and assembly remain uncharacterized for the most part of proteomes. However, previous work has shown that static structural data can be leveraged in computational analyses to gain insights into protein folding ^4,5,14^ and assembly mechanisms ^15–20^. This idea motivates us to leverage the wealth of structural data to gain insights into molecular determinants of the dynamic process that is co-translational assembly.

A key distinction in the assembly pathway of protein complexes lies in the post-or co-translational timing of assembly (**Fig. 1A**). In the latter, translation, folding, and assembly occur simultaneously, which can help funneling the assembly pathway of complexes, minimizing promiscuous interactions, or regulating orphan subunit degradation ^21,22^. Recent reports have shown that co-translational assembly is prevalent ^23–27^. In particular, Bertolini et al. ^28^ identified thousands of proteins undergoing co-translational assembly by sequencing footprints protected among isolated disomes, *i*.*e*., pairs of ribosomes connected by their respective, interacting, nascent chains. Because disome footprints are sequenced over an entire lysate, such an experiment can provide a list of co-translationally assembling protein subunits – here referred to as ‘coco’ subunits – but does not inform on interacting partners. These limitations motivated us to integrate these disome data with protein structures, both to uncover specific molecular signatures of co-translational assembly as well as identify specific interacting pairs undergoing this process in cells.

**Fig. 1.**
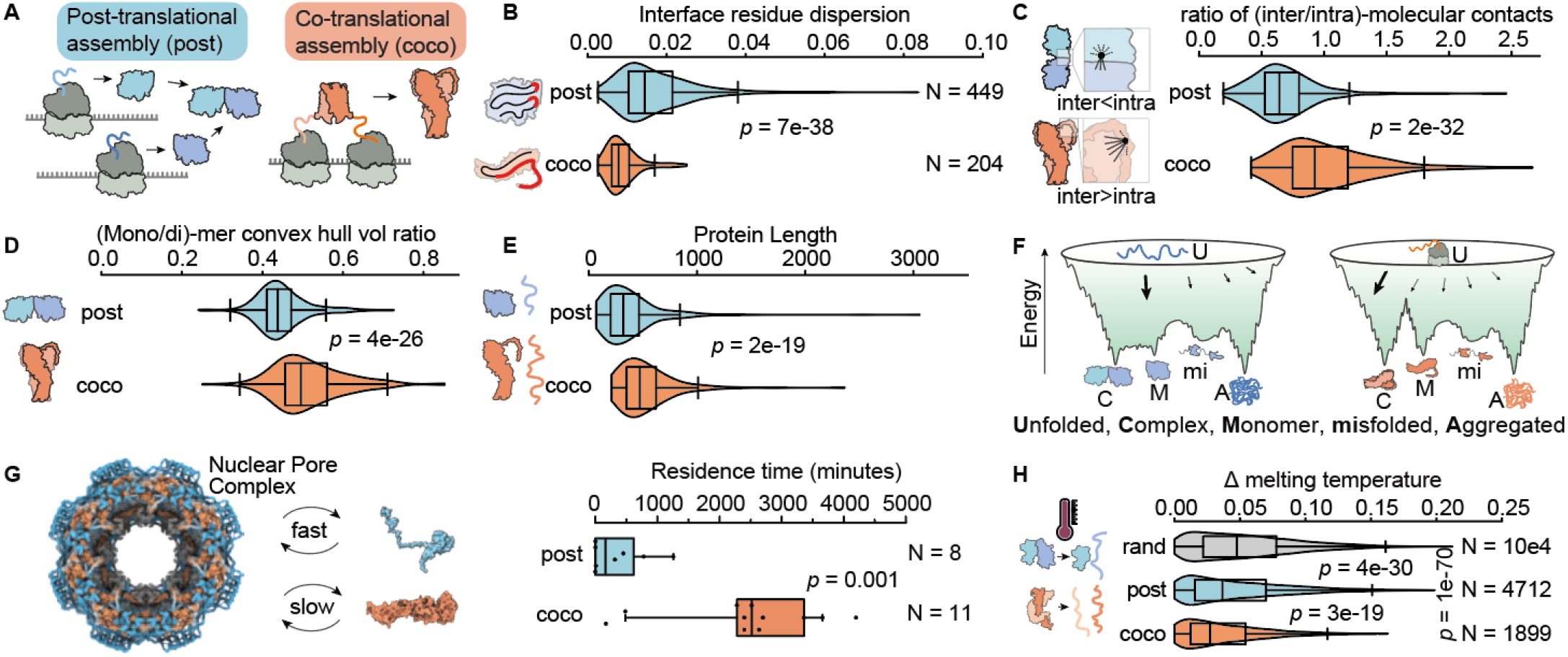
Subunit interaction geometry underlies co-or post-translational modes of assembly. (**A**) Subunits are classically assumed to be translated and to fold independently before they interact (left, cyan). By contrast, in co-translational assembly (right, orange), subunits start interacting during their translation. (**B**) Violin plots comparing the dispersion of interface residues in the sequence among co-(orange) and post-translational (cyan) homomers of known structure. The boxes and whiskers indicate the 25^th^-75^th^ and 5^th^-95^th^ percentiles of the distributions. The central line shows the median. Differences between distributions are assessed by the Mann-Whitney U test. (**C**) Same as (B) for the ratios of inter-to intra-molecular atomic contacts of interface residues. (**D**) Same as (B) for the ratio of convex hull volume enclosing the monomer relative to that enclosing the dimer. (**E**) Same as (B) for the protein length. (**F**) Hypothetical energy landscapes associated with post- and co-translational assembly. The intertwined structure of coco-subunits implies that extensive conformational rearrangements are required for free monomers to bind to each other. Such rearrangements may be associated with a high activation energy, which co-translational assembly could circumvent by shunting the unbound, monomeric state. **(G)** We analyzed human nuclear pore complex (PDB code 7R5K ^34^) subunit exchange kinetics measured *in vivo* ^35^. We assigned subunits to the post (cyan) or coco (orange) category based on disome enrichment data. The boxplot shows residence time distributions for coco and post subunits. The coco subunits show slower exchange kinetics, in line with a high energy barrier separating their unbound and bound states. **(H)** Same as (B) for melting temperature deviations of random protein pairs (gray), post-(cyan), and coco-pairs (orange) from heterodimers. The procedure to assign heterodimers as post-or coco pairs is described in *Methods 3*.

Our analysis uncovered that co-translational assembly is governed by structural and biophysical features of protein complexes and often involves intertwined and mutually stabilized subunits. This structural feature implies that complexes assembling co-translationally are more kinetically stable than complexes assembling post-translationally, which is manifested in longer subunit residence times and similar thermal denaturation temperatures of interacting subunits. Additionally, we find that the inter-dependence of interacting coco subunits necessitates their co-regulation, presumably to balance their stoichiometry. Utilizing these concepts, we developed a model relying on structural data to predict co-or post-translational assembly in both homo- and heteromeric complexes. We validated these predictions through established disome footprint data and targeted experiments. These findings pave the way to a better understanding of protein folding and assembly beyond the textbook knowledge of reversible folding and binding, and they highlight the overarching impact of protein structure on translation and expression regulation.

## RESULTS

### Subunit interaction geometry underlies co- and post-translational modes of assembly

We mapped the comprehensive dataset of human coco subunits from reference ^28^ onto available 3D structures of homo- and hetero-oligomeric complexes ^29,30^. We anticipated coco subunits to show distinct structural characteristics, notably intertwined subunits, which may suggest a concurrent folding-binding mechanism ^31–33^. Indeed, intertwined subunits, stabilized through extensive contacts with their partner, are expected to be unstable as monomers and so could use co-translational assembly as a means to bypass an unstable monomeric state. To capture relevant geometric and structural features, we employed a combination of established and newly defined metrics (*Methods 1-2*).

We analyzed such signatures in coco and ‘post’ (referring to post-translationally assembling) subunits, using protein complexes of known structure. We initially focused on homo-oligomers as they alleviate the need to infer coco subunit pairs, which are not known. We identified 204 and 449 homo-oligomers of known structure experimentally classified as coco or post-subunits respectively.

Several structural properties of these complexes showed significant differences between the two groups (**Fig. 1 B-E**). Notably, interface residues appeared more clustered in the sequence among coco subunits than among post (**Fig. 1B**, Mann-Whitney U-test *p* = 7E−38). Such localized interaction sites may decrease the necessity to synthesize and fold long segments before binding, thereby promoting co-translational partner recognition. In a different analysis, we found the interface residues exhibited a higher ratio of inter/intra molecular contacts among coco subunits relative to post (MWU *p* = 2E−32, **Fig. 1C**). The pronounced inter-molecular contacts imply that coco subunits are equally stabilized by their partner (inter) as they are by their own tertiary structure (intra). Such partner-dependent stability is reminiscent of domain-swapping, which we analyzed through the *V*_*m*_/*V*_*d*_ ratio, of convex-hull volumes enclosing the monomer (*V*_*m*_) and dimer (*V*_*d*_). Indeed, the larger ratios observed for coco-subunits relative to post (MWU *p* = 4E−26, **Fig. 1D**, *Methods 2*.*5*) indicate pronounced intertwined structures. Interestingly, large interface areas have previously been linked to co-translational assembly ^32^, and our findings suggest that the underlying determinant is the subunit interaction geometry (*Supplementary Note 1-2*). Finally, we observed that coco subunits were encoded by longer polypeptide chains than post-subunits (MWU *p* = 2E−19, **Fig. 1E**), presumably promoting binding while the C-terminal region is still being synthesized.

These geometrical features, along with others shown in *Supplementary Note 1*, reflect a proto-typical coco-subunit as binding its partner through a local region in the sequence. Moreover, this binding region typically extends into the partner’s structure, intertwines with it, and is thereby partner-stabilized. Such geometrical features imply that conformational rearrangements must occur upon binding, and we expect such rearrangements to create a high energy barrier that co-translational folding-binding helps overcome (**Fig. 1F**) ^33,36,37^. In that respect, we predict that complexes assembled co-translationally are more kinetically stable than those assembled post-translationally.

We investigated whether geometrical and kinetic observations were reproduced among heteromeric complexes. To this aim, we projected the disome enrichment data onto complexes of known structure (*Methods 2*). The differences in structural features were recapitulated among hetero-oligomers as well (*Supplementary Note 1*). To assess differences in kinetic stability, we analyzed the residence times of subunits in the human nuclear pore complex, as previously measured *in vivo* by fluorescent labeling ^35^. Consistent with expectations, residence times of coco subunits were about five-fold longer than post subunits (**Fig. 1G**, MWU *p* = 0.001). Additionally, we hypothesized that coco protein pairs should melt at similar temperatures upon thermal denaturation. We analyzed a comprehensive dataset of *T*_*m*_ values measured for human proteins ^38^, revealing that coco subunits exhibit similar *T*_*m*_ values (low Δ*T*_*m*_) when compared to post-subunits (coco < post < random, MWU *p* < 3E−19, **Fig. 1H**, also see *Supplementary Note 3*).

Taken together, these observations highlight that specific structural features introducing partner-dependent stability characterize coco subunits.

### Proteostasis of coco subunit pairs is synchronized across the central dogma

The structural interdependence of coco subunit pairs suggests that their genes should be expressed together and that their mRNAs should be translated synchronously to yield 1:1 stoichiometry at the protein level. To assess such synchrony, we compared the similarity of expression and regulation between heteromeric coco and post pairs, across the central dogma (**Fig. 2A**). To this aim, we projected the disome enrichment data onto a dataset of human heterodimers integrated from multiple repositories (*Methods 3-4*), resulting in 19,855 pairs of which 4,016 and 15,839 were assigned as coco and post pairs respectively.

**Fig. 2.**
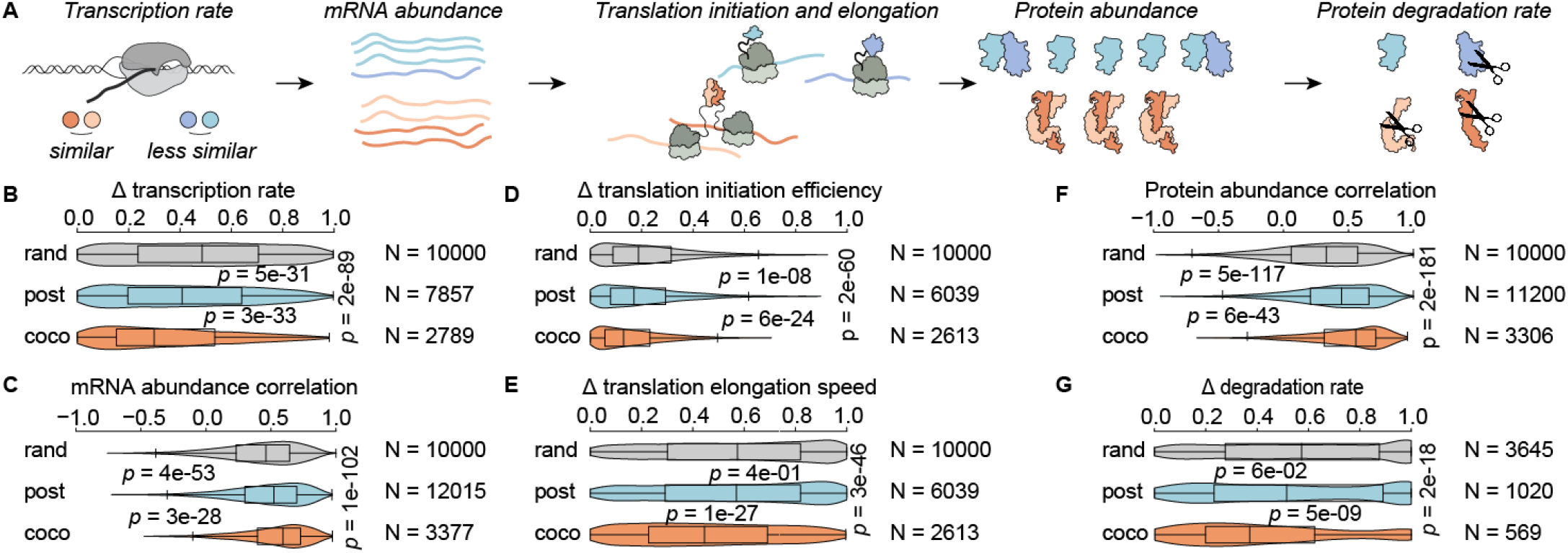
Co-translational heteromeric pairs undergo synchronized regulation across the central dogma. (**A**) Schematic representation of the central dogma. (**B**) Violin plots comparing transcription rate deviations of co-(orange) and post-translational (cyan) heteromeric pairs. Random pairs are shown for reference (gray). Figure features follow Fig. 1B. (**C**) Similarity in mRNA abundance of coco or post subunit pairs measured as the Spearman correlation across 273 RNA-seq datasets from Bgee ^40^. (**D**) Same as (B) for mRNA translational initiation efficiency. (**E**) Same as (B) for mRNA translation elongation speed. (**F**) Same as (C) for protein abundance compiled from 156 proteomic datasets from PaxDB ^41^. (**G**) Same as (B) for protein degradation rates.

We initially analyzed transcription rates measured in human cells ^39^, revealing that coco pairs show more similar rates than post pairs (Δ transcription rate of coco < post < random, MWU *p* < 5E−31, **Fig. 2B**). Consistent with this observation, mRNA abundance levels from 273 RNA-seq experiments covering 53 human tissues ^40^ showed higher correlations for coco pairs than for post pairs (correlation of coco > post > random, MWU *p* < 3E−28, **Fig. 2C**).

At the translational level, we analyzed ribosome profiling experiments ^42^. Translation appeared more synchronized in coco pairs than in post, with the former exhibiting more similar initiation (Δ translation initiation efficiency of coco < post < random, MWU *p* < 6E−24, **Fig. 2D**) and elongation rates (Δ elongation rate of coco < post < random, MWU *p* < 1E−27, **Fig. 2E**). Furthermore, the variation of protein abundance measured across 153 datasets ^41^ showed higher correlation among coco subunits than among post, reflecting more consistent stoichiometries in the former type (correlation of coco > post > random, MWU *p* < 6E−43, **Fig. 2E**).

Beyond translation, we expected the partner-dependence for stability among coco pairs to be reflected in synchronized degradation of these subunits. Indeed, a dataset of degradation rates characterized for human proteins ^43^, revealed more similar rates for coco pairs than for post (Δ degradation rate of coco < post < random, MWU *p* < 5E−9, **Fig. 2F**).

Taken together, these results depict how subunit interaction geometry underlies the mode of complex assembly, which in turn shapes the global regulatory landscape of the respective protein subunits.

### Predicting co-translational assembly on a proteome-wide scale from structure

The marked geometric differences in the structures of coco and post subunits, prompted us to predict the mode of assembly of a complex solely based on its structure. We initially focused on homo-oligomers as these alleviate the need to infer coco pairs from the experimental disome enrichment data. Since Badonyi and Marsh ^32^ previously used Bertolini et al.’s ^28^ coco/post definition to associate large interface areas with co-translational assembly, we asked how well a prediction model performs on their existing data, and whether our analyses would make any significant improvements.

We fitted a logistic regression model using Interface area for Badonyi and Marsh’s dataset of 1351 homomeric structures ^32^, and assessed the prediction accuracy using a receiver operating characteristic (ROC) curve. Interface area recapitulated the experimental characterization of coco/post homomers with an area under the curve (AUC) of 0.629 (brown, **Fig. 3A**). Similar results were obtained when our structure dataset and coco/post definitions were used instead (green, AUC=0.751). Other orthogonal parameters such as ratio of inter/intra-molecular contacts captured the experimental data with a relatively higher accuracy (black, AUC=0.795), but no single parameter achieved highly accurate predictions (*Supplementary Note 2*). Remarkably, a dramatic improvement was obtained when eight structural features (each summarized as a single number) were integrated to capture the mode of assembly (AUC=0.940, Average Precision, AP=0.901, *Methods 5*). These highly accurate predictions motivated us to scale them up to entire proteomes.

**Fig. 3.**
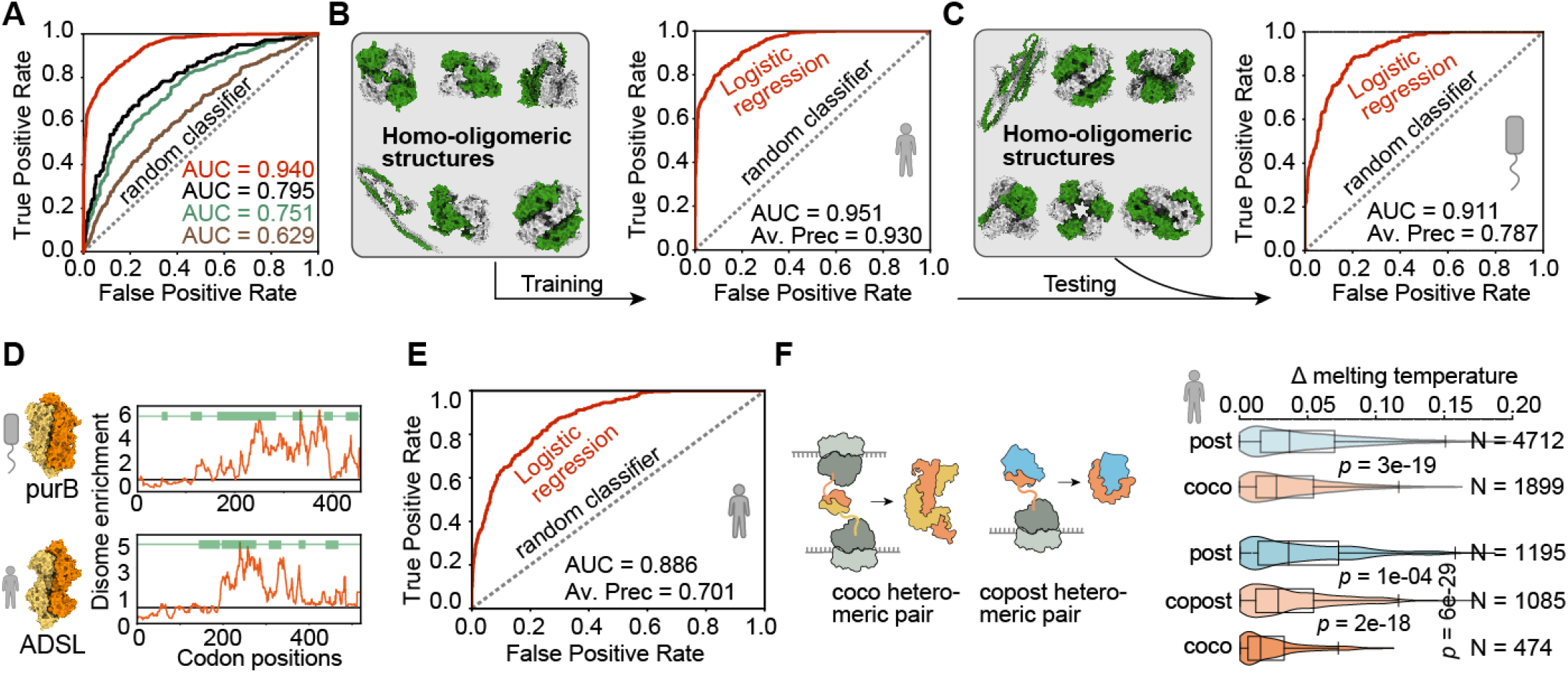
Protein structure captures proteome-wide information on co- and post-translational assembly. (**A**) The interface area is larger among coco complexes compared to post, but this criteria alone showed limited predictive power in a previous work (AUC=0.629, brown, (30)) as well as in our dataset (AUC=0.751, green). Other orthogonal structural features (e.g., ratio of inter/intra-molecular contacts) can capture the experimental data with higher accuracy (AUC=0.795, black), but no single parameter achieves high-accuracy predictions (Supplementary Note 2). Remarkably a logistic regression model utilizing eight structural features, each summarized as a single number, achieves high accuracy predictions (AUC=0.940, red). (**B**) Extending the dataset of 653 experimental structures used in panel A with 1395 homo-oligomeric structures predicted with AlphaFold2 ^44^ (*Methods 2*.*1* and *2*.*2*) improves both the coverage and accuracy of our predictions using the same eight-parameter model (**C**) Same as (B), for the bacterium *Escherichia coli*. Prediction results of the model trained on human structures, now predicting the assembly mode of *E. coli* homomers. (**D**) The conservation of structure and co-translational assembly is depicted for two orthologous enzymes: *E. coli* purB and human ADSL (PDB codes: 2PTQ ^49^ and 4FFX ^50^). We show the disome footprint enrichment relative to monosomes (red), smoothened over a 15-codon sliding window. Values above the line (y=1) represent a disome enrichment and thereby co-translational assembly. Interface residues are highlighted with green rectangles. (**E**) Same as (B), the ROC curve for predicting the co- and post-translational modes of heteromer assembly (*Methods 5*). **(F)** Co-translational heteromer assembly can be bidirectional (coco) or unidirectional (copost). Transparent violins replicate **Fig. 1H** and are compared to predictions after incorporating the uni-or bi-directionality of co-translational assembly based on structure.

Recent advances in machine learning have increased the coverage of protein tertiary structures ^44^ and protein complexes ^45–48^ to entire proteomes. We integrated an atlas of 2012 homodimers (**Fig. 3A**, *Methods 5*), of which 672 and 1376 were assigned as coco and post respectively based on the experimental data. We fitted a logistic regression model integrating eight topological features, which together captured the mode of assembly accurately (AUC = 0.951; **Fig. 3B**; *Supplementary Note 2*). To assess the generality of this model, we retrieved disome-enrichment data for *E. coli* ^28^, which served as a test set. Interestingly, these data were indirectly identified as part of an experiment analyzing disomes formed by a human protein heterologously expressed in *E. coli*, and were not analyzed before to describe complexes from *E. coli*. We mapped these disome-enrichment data to homo-oligomeric structures of *E. coli* and inferred a coco probability for each complex, based on the model trained on human proteins. These predictions recapitulated well the experimental disome enrichment (AUC = 0.911; **Fig. 3C**), validating our model and its underlying assumption that co-translational assembly is primarily determined by the structure of a complex.

A remarkable evolutionary implication of our findings is that co-translational assembly should be conserved in evolution because protein structure is conserved itself. We provide such an example with two orthologous enzymes involved in purine biosynthesis: purB (*E. coli)* and ADSL (human). Their translation shifts from monosomes to disomes near the 200^th^ codon, coinciding with the emergence of interface residues at the ribosome tunnel (**Fig. 3E**).

Accurately predicting homodimers’ assembly mode from structure motivated us to apply the same strategy to heterodimers. While disome enrichment experiments do not provide specific interaction partners, we mapped putative coco and post partners in a two-step process (*Methods 5*). Briefly, we first selected interacting protein pairs of known structure with both subunits identified as either coco or post. We fitted a first predictive model based on this set using the same topological features, and kept only the highest-scoring complex for each protein. The rationale for this selection is that a protein may engage in multiple post-interactions (low scores), and we assumed there should be fewer (typically a single) co-translational interactions. The resulting set consisted of 679 and 2113 heterodimer structures assigned as coco and post respectively. Refitting our structural model onto this filtered set recapitulated the experimental data well (AUC = 0.886, **Fig. 3F**). Additionally, this model also generalized to heterodimers from *E. coli*, recapitulating these independent experimental data with an AUC value of 0.841 (*Supplementary Note 2*).

Remarkably, because some of the structural features are directional, this model provides two distinct probabilities per complex, *i*.*e*., one for each subunit. We applied this model to 7299 protein pairs of known structure, which we could now classify as post (3945 pairs, low score for both partners), coco (1246 pairs, high score for both partners), but also co-post (2108 pairs, only one partner with a high score). The latter type points to proteins that bind co-translationally to fully-translated partners.

As an additional validation of the model’s predictive value, we examined whether it reproduced observations based on experimental data, specifically where coco pairs exhibited similar melting temperatures or co-regulation across the central dogma. Notably, the similarity in melting temperatures between proteins in the coco pairs predicted based on structure was even greater than that observed in coco pairs defined based on disome-enrichment data (**Fig. 3G**). Consistent with this observation, coco pairs predicted based on structure also showed greater synchrony across the central dogma than coco pairs defined based on experimental data (*Supplementary Note 4*).

These results show that co-translational assembly can be accurately predicted from structure, and imply that coco subunits and pairs can be inferred from any proteome for which structural information is available. This idea motivated us to gain a comprehensive overview of co-translational assembly across the *H. sapiens, E. coli*, and *S. cerevisiae* proteomes, although the latter had no systematic experimental information on co-translational assembly.

### A comprehensive view of co-translational assembly from structure

We integrated structure-based predictions with experimental disome-enrichment data to gain an overview of co-translational assembly across the human proteome. In this integration, structure-based predictions added 561 proteins annotated as coco, yielding a total of 3073. Thus, about 17% of the proteome was estimated to assemble co-translationally, about 53% was annotated as post, and the remaining ∼31% could not be annotated due to ambiguous or absent data (**Fig. 4A**). Focusing our analysis on 1616 heteromeric complexes of known structure, we predicted that 23%, 14%, and 63% of subunits assemble in coco, copost, and post manners, respectively. We reached similar conclusions for *E. coli*, where the proteome fraction of coco subunits was estimated at ∼20% (**Fig. 4A**).

**Fig. 4.**
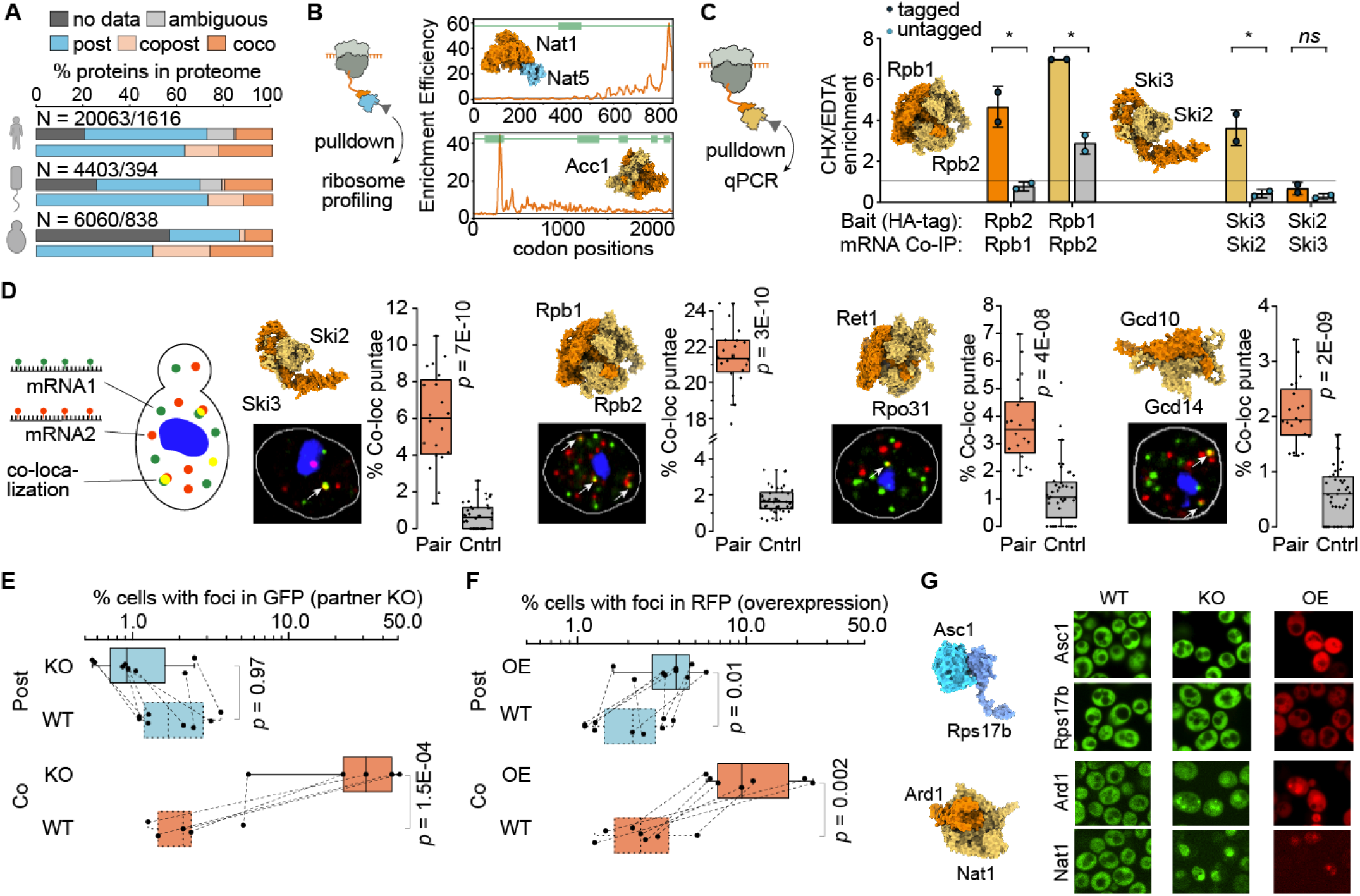
De novo prediction of co-translational assembly across the yeast proteome. **(A)** Global statistics on co/post-translational assembly for the proteomes of human, *E. coli, and S. cerevisiae*. For each organism, the top and bottom bars show statistics for the reference proteome and for heteromeric complexes of known structure. (**B**) Quantifying co-translational assembly of Nat1/5 heteromer (top, PDB code: 6O07 ^52^) and Acc1 homodimer (bottom, PDB code: 5CSL ^53^) by SeRP. The average ribosome footprint enrichment of two biological replicates is shown as an orange line. Interface residue positions are highlighted by green boxes. (**C**) Quantifying co-translational assembly by RIP-qPCR for Rpb1/2 (PDB code: 1Y1V, ^54^) and Ski2/3 (PDB code: 4BUJ ^55^). Barplots depict mRNA enrichment after pull-down following cycloheximide (CHX) *vs*. EDTA treatment. We show the enrichment after pull down of a HA-tagged (orange/yellow bars) or untagged (negative control, gray bars) protein partner. A horizontal line (y = 1) represents no enrichment; error bars show the standard deviation of two biological replicates. Statistical significance was assessed by two-sample t-tests: ns *p* > 0.05, ^*^ *p* < 0.05. (**D**) RNA-smFISH experiments in yeast cells reveal specific co-localization of mRNAs for four predicted coco pairs: Ski2/3, Rpb1/2, Ret1/Rpo31 (PDB code: 5M5W ^56^), and Gcd10/14 (ModelArchive code: ma-bak-cepc-0793 ^46^). Fluorescent tags: blue, DAPI (nucleus); red, CAL Fluor Red 590; green, Quasar 670. Co-localizing spots are yellow. The box plots depict the distribution of percent co-localized punctae per field of view; negative controls consist of two mRNAs encoding unrelated proteins (*Methods 8, Supplementary Note 7*). Statistical significance was assessed by the MWU test. (**E**) We evaluated the tendency of coco (orange) or post (blue) subunits to become insoluble and form punctae upon partner knockout or (**F**) over-expression. In both panels, paired boxplots show the percentage of cells containing one or more punctae upon deletion of a partner (KO) or upon over-expressing the protein (OE); *p*-values are calculated using the MWU test. (**G**) Example structures of two candidate pairs (PDB codes: 6HD5 ^57^, 7A1G ^58^) and corresponding images in KO or OE experiments.

Although no disome enrichment information is available for the proteome of *S. cerevisiae*, we integrated structural information from the Protein Data Bank ^30^ as well as AlphaFold2-based structures for homo-^45^ and hetero-oligomers ^46^. These structural data enabled us to derive proteome-wide predictions (**Fig. 4A**) that we subsequently evaluated with targeted experiments.

We first evaluated the predictions using selective ribosome profiling (SeRP) ^24^ We compared ribosome footprints from the total translatome to those co-purified with a C-terminally tagged protein (*Methods 6*). Thus, co-purified footprints point to a co-translationally interacting partner. We employed SeRP to analyze the Nat1/5 heterodimer (NatE complex) predicted as copost with translating Nat1 assembling onto full-length Nat5. Upon isolating tagged Nat5 we indeed co-purified ribosomes translating Nat1 mRNA. The footprint density of Nat1 mRNA increased onwards of codon 480, coinciding with the ribosomal exit and exposure of its binding interface (**Fig. 4B**). Next we examined the Acc1 (Acetyl-CoA carboxylase) homodimer, also predicted to assemble co-translationally. SeRP uncovered that C-terminally tagged full-length Acc1 similarly co-purifies with ribosomes translating its own mRNA, and the increased footprint density onwards of codon 250 coincides with the ribosomal exit of its interface (**Fig. 4B**, also see *Supplementary Note 5*).

We next utilized RNA immunoprecipitation qPCR (RIP-qPCR) ^51^ whereby translating mRNAs physically associated with the C-terminally tagged partner are quantified by qPCR after pull-down (*Methods 7*). Although RIP-qPCR primarily detects copost assembly, it could in principle also capture coco events when the tagged subunit is released by the ribosome before its partner (*e*.*g*., due to C-terminus proximity of the interface). We first examined a predicted coco pair Rpb1/2, forming the core of the RNA polymerase-II complex. We pulled down on tagged Rpb1, in conditions promoting ribosome stability (cycloheximide) or dissociation (EDTA), and quantified Rpb2 mRNA levels. We observed an enrichment in Rpb2 *mRNA* that was cycloheximide-dependent and also tag-dependent, indicating that Rpb2 binds Rpb1 co-translationally. These observations were mirrored when pulling down on tagged Rpb2, suggesting that this pair assembles co-translationally (**Fig. 4C**).

We subsequently examined the Ski2/3 dimer of the RNA-helicase Ski complex, also predicted as coco. RIP-qPCR experiments indicated that translating Ski2 assembled with full-length Ski3, but the reverse interaction was not observed (**Fig. 4C**). This discrepancy may originate in the interface of Ski3 being closer to the C-terminus than Ski2 (*Supplementary Note 6*). Therefore, the release of the Ski2 C-terminus and its pull-down are unlikely to occur while Ski3 is being translated. A similar asymmetry was observed for the Rrn6/7 pair of the RNA polymerase-I core factor complex (*Supplementary Note 6*).

We reasoned that coco assembly is expected to yield co-localized mRNAs, motivating us to characterize the Ski2/3 complex further by single-molecule RNA-fluorescence in situ hybridization (smFISH) ^59^. Such mRNA co-localization has rarely been observed to our knowledge ^51,60^, perhaps due to the difficulty in reliably identifying coco subunit pairs. We designed FISH probes conjugated to two different colors, each specific to the mRNA encoding the subunit of a predicted coco pair. After hybridization, we visualized the subcellular localization of each mRNA by fluorescence microscopy and quantified the co-localization of each pair (*Methods 8, Supplementary Note 7*). The mRNAs encoding Ski2 *and* Ski3 showed significant co-localization when compared to two unrelated genes (MWU *p* = 7E−10, **Fig. 4D**), consistent with their predicted bidirectional coco assembly. We observed similar results for Rpb1/2, Ret1/Rpo31 (RNA polymerase-III complex), and Gcd10/14 (tRNA methyltransferase complex). Interestingly, the four pairs showed various extent of co-localization, from 2% to 21% (**Fig. 4D**), which may be explained by several factors that include the fraction of actively translated mRNAs, mRNA hybridization and detection efficiency, and potential alternative assembly pathways not involving co-translational assembly. Remarkably, the Gcd10/14 pair was predicted based on an AlphaFold2-predicted structure ^46^.

The structural and omics analyses from Figs. 1 and 2 show that coco subunits show partner-dependent stability and solubility, while post-subunits appear more independent. We examined such partner-dependence among yeast complexes predicted as coco and post. We tagged both types of subunits with a fluorescent protein and assessed their intracellular solubility based on the formation of puncta by fluorescence microscopy (*Methods 9*). We explored two conditions that interfere with 1:1 stoichiometry, namely partner knockout or overexpression. We found that coco subunits were more prone to form puncta than post subunits, both when their partner was knocked out, or when they were over-expressed themselves (**Fig. 4E-G**). Thus, both types of experiments suggest that dosage balance and stoichiometric expression of coco subunits are critical for proper cellular functions, an observation consistent with the synchrony seen across the proteostasis steps (**Fig. 2**).

Finally, we analyzed a shotgun proteomics dataset of thousands of knock-out strains ^61^, which supported these observations further: while the abundance of proteins predicted to assemble post-translationally did not change significantly in the absence of their partner, the abundance of coco subunits dropped significantly. Remarkably, for copost pairs, the drop in protein abundance was consistent with the predicted direction of assembly (*Supplementary Note 8*).

## DISCUSSION AND CONCLUSIONS

We elucidated structural features facilitating co-translational assembly, including clustered interface residues, partner-stabilized intertwined interfaces, and prolonged translation, as reflected in longer protein sequence lengths. Additionally, coco subunits appeared structurally unstable as monomers and indeed, showed reduced solubility upon changes in partner stoichiometry. Accordingly, we saw that regulatory mechanisms ensure balanced stoichiometries in co-translational pairs, including synchronized transcription, translation, and degradation. In contrast, proteins that assemble post-translationally were typically stable, as indicated by their globular structures and their maintained solubility upon changes in partner stoichiometry.

Interestingly, the connection between co-translational assembly and structure suggests that two protein partners can start binding during synthesis if their emerging chains are in each other’s vicinity. When considering homo-oligomers, this property is naturally met by polysomes. In this context, our results corroborate that biasing interface residues towards the C-terminus can ensure minimal interference between folding and binding ^62^. Considering heteromers, mRNA co-localization is a prerequisite for co-translational assembly. Such co-localization could be achieved actively through zipcode sequences ^63^ or through partitioning into specific condensates ^64^. Alternatively, random diffusion could suffice to promote nascent chain recognition ^65^. Remarkably, the coco partners we predicted from structure showed marked mRNA co-localization as seen by smFISH, and thereby constitute a rich dataset to explore these alternative hypotheses.

Equally important, the conservation of structure over evolutionary timescales implies co-translational assembly to be conserved as well. In that respect, our predictive model and proteome-wide datasets offer solid grounds on which to base future comparative analyses of protein assembly.

We unveiled a fundamental connection between protein structure and the translation process, emphasizing its impact on gene expression, proteostasis, and mRNA localization. The resulting comprehensive description of protein co-translational assembly will serve as a valuable framework for future studies in protein interaction, regulation, and evolutionary biology, promoting a holistic understanding of proteome organization and functionality.

## Acknowledgments

We thank Amnon Horovitz, Jeffrey Gerst, Yitzhak Pilpel, Daniel Dar, Sarah Teichmann, Judith Frydman, Bernd Bukau, Günter Kramer, Stephen Fried, Shimon Bershtein, Nir Fluman, Sudip Kundu, and all members of E.D.L and A.S.’s lab for helpful discussions; Jeffrey Gerst, Shahar Garin, and Gal Haimovich for advice and reagents on smFISH; Liat Alyagor and Roni Oren for advice on smFISH and help with testing different microscopes.

## Author contributions

S.M., A.S., and E.D.L. conceived and conceptualized the research; S.M. performed the structural and OMICs analysis shown in Figs. 1 and 2, and co-translational assembly predictions shown in Figs. 3 and 4A; J.V. performed the SeRP and RIP-qPCR experiments shown in Fig. 4B, 4C and Supplementary Note 5; A.L. performed the smFISH experiments shown in Fig. 4D and Supplementary Note 6; M.H. and H.G.S. performed the genetic perturbation experiments shown in Fig. 4E;A.S. supervised the SeRP and RIP-qPCR experiments; T.O.Y. conceived and guided S.M. with structural analyses on ‘rigid body entanglement’ and ‘convex hull’; E.D.L. supervised the computational analyses, smFISH, and genetic perturbation experiments; S.M. prepared all the figures; E.D.L. edited the figures; S.M. and E.D.L. wrote the paper; A.S. and E.D.L. acquired funding; all authors discussed and commented on the results and edited versions of the paper.

## Data and materials availability

All predictions and datasets are provided as Supplementary Material or are available on FigShare (https://doi.org/10.6084/m9.figshare.24311917.v1).

## Code availability

Code and custom scripts used in this work are available from the authors upon request.

