## Supplementary Methods for "Structural determinants of co-translational protein complex assembly"

### The structure of protein complexes underlies co-translational assembly

**This Supplementary file includes:**

Supplementary Methods Sections 1-9
Supplementary Figures S1 to S6
Supplementary Table S1
Supplementary References

**Supplementary Datasets for this manuscript include the following:**

**Data S1**. Structure Dataset for *Homo sapiens*
**Data S2**. Structure Dataset for *Escherichia coli*
**Data S3**. Structure Dataset for *Saccharomyces cerevisiae*
**Data S4**. Human co-complex subunit pairs
**Data S5**. Delta-Melting-Temperature matrix for human
**Data S6**. Melting-Curve-Dissimilarity matrix for human
**Data S7**. Protein-Abundance-correlation matrix for human
**Data S8**. Delta-Protein-Degradation-Rate matrix for human
**Data S9**. Delta-Transcription-Rate matrix for human
**Data S10**. Gene-Expression-correlation matrix for human
**Data S11**. Delta-Translation-Initiation-Efficiency matrix for human
**Data S12**. Delta-Translation-Elongation-Speed matrix for human
**Data S13**. Fluorescent probe sequences used for smFISH experiments
**Data S14**. Plasmids generated for genetic perturbation experiments
**Data S15**. Yeast strains generated for genetic perturbation experiments

Supplementary data files are available in FigShare (DOI: 10.6084/m9.figshare.24311917).

### Materials and Methods

#### 1. Deriving high-confidence sets of co- and post-translationally assembling proteins from disome-enrichment data

The initial dataset of 15980 human proteins with characterized disome over monosome enrichment was obtained from Bertolini et al. (*1*). The authors distinguished coco subunits based on the messenger RNA (mRNA) enrichment in the disome fraction relative to the monosome fraction, as inferred from sequencing reads of the ribosome footprints. Indeed, as nascent chains emerge and begin to interact, the density of disome footprints associated with their mRNAs is expected to increase relative to footprints identified in monosomes for the same mRNAs. In the original work (*1*), disome enrichment was calculated using footprints spanning the full-length mRNA. Here, we used the same measure, and added an additional parameter reflecting the locality or globality of the enrichment across the mRNA. Upon inspection of the profiles, we noticed that some proteins were characterized by a localized peak enrichment (**Fig. S1A**), whereas others showed an enrichment throughout a wider region (**Fig. S1B**). We reasoned that co-translational assembly should be associated with the latter type. To distinguish both types, we analyzed the average disome enrichment as a function of the peak enrichment reflected in the 95th percentile value of the profile (**Fig. S1C**). In this representation we defined a line $y = 3.7x - 1.65$ (green dotted line), representing proteins with a peak enrichment equal to 3.7-fold the mean enrichment minus a constant (1.65). Points above this line have a stronger peak-to-mean enrichment than proteins below this line. Accordingly we defined coco subunits as those below this line and also showing a mean value ≥1.50 (**Fig. S1B**, gray region). By contrast, we considered as post-subunits those above the line $y = 3.7x - 1.25$, or with a mean enrichment below 1 (blue region). Finally, the proteins situated in between these two regions were classified as ambiguous (yellow region).

In this manner we defined 2512, 8547, and 3392 proteins as being coco, post, or ambiguous subunits.

The work of Bertolini et al. includes an experiment of human genes over-expressed in *E. coli* to assess their co-translational assembly upon heterologous expression. Interestingly, these experiments characterized disome-enrichments for the human genes and alongside identified those mediated by *E. coli* proteins as well. We analyzed this dataset using the same methodology, which resulted in a set of 783, 1833, and 504 coco post, and ambiguous subunits in *E. coli* (**Fig. S1D**).

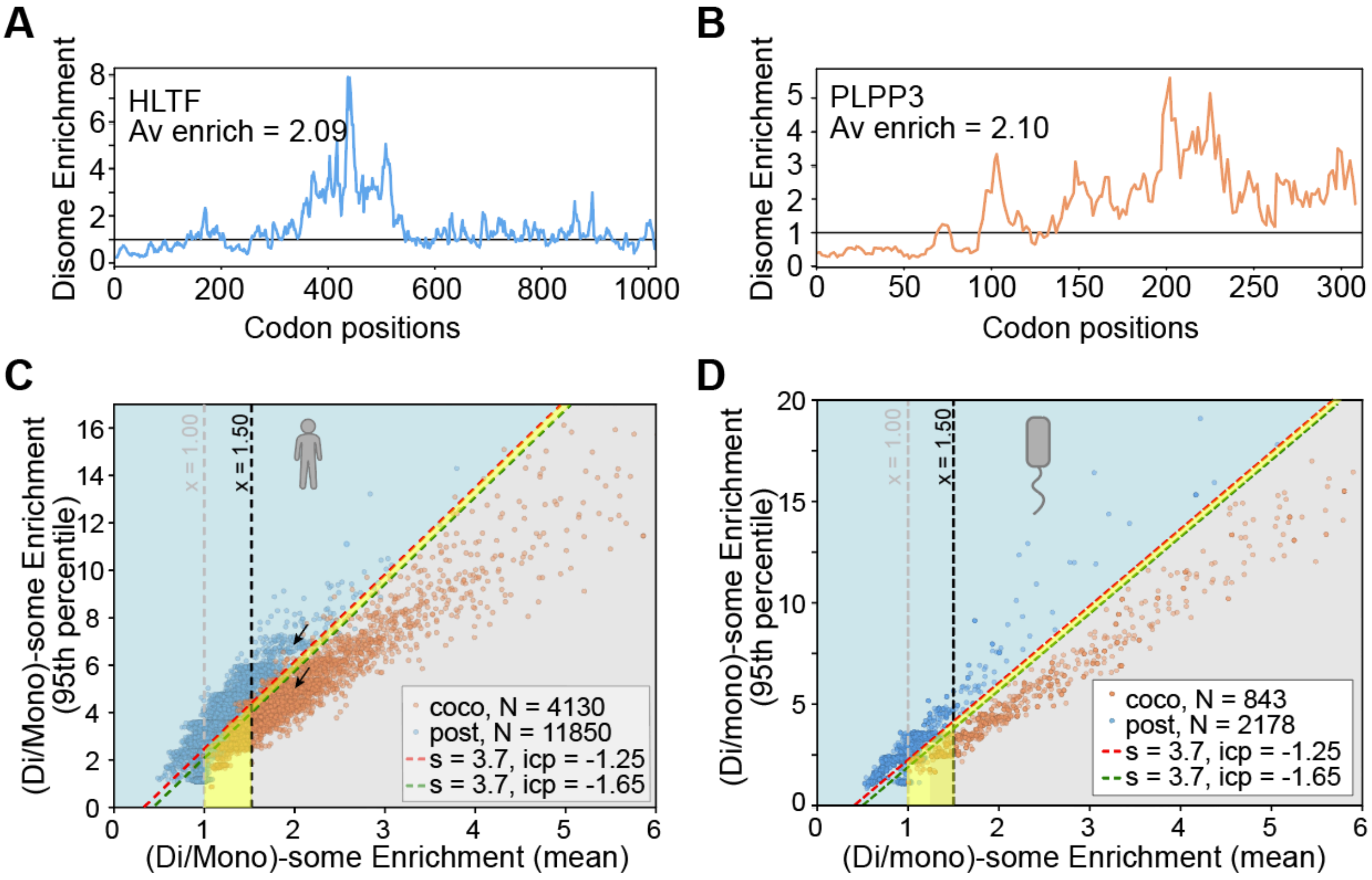


**Fig. S1**. (**A**) The disome enrichment profile of human HLTF gene, window averaged over 15-codon. **(B)** Same, for the PLPP3 gene. Both HLTF and PLPP3 exhibit similar mean disome enrichments (2.09 and 2.10 respectively). However, the enrichment profile of HLTF is localized around codons 350-550. In contrast, PLPP3 shows a more persistent enrichment after the onset around codon 90, until the end of translation. (**C**) We expect coco subunits to remain bound during translation, after their initial encounter. We assessed such enrichment persistence along the mRNA by the peak-enrichment (y-axis) relative to the mean-enrichment (x-axis). We capture such “persistence” in a scatter plot showing the mean (x-axis) versus 95th percentile (y-axis) disome enrichment, for 15980 human genes. Vertical dotted lines highlight mean enrichment thresholds equal to 1.0 and 1.5. The red and green dotted lines have the same slope (s) and different intercept (icp) values, and segregate well the coco and post subunits as defined by Bertolini et al. (*2*). Here, we take the same definition of coco- and post subunits with an additional constraint based on this representation: coco-type subunits are those previously identified that also lie in the gray region (mean enrichment above 1.5, and peak enrichment not exceeding 3.7*mean − 1.65). By contrast, post subunits are those in the blue region (mean enrichment below 1, or peak enrichment exceeding 3.7*mean − 1.25). Finally, all the points not matching these criteria were assigned as ambiguous in terms of their co-translational assembly status (yellow region as well as coco-subunits in the blue region and post-subunits in the gray region). (**D**) Same data as panel (B) for *Escherichia coli*.

#### 2. Structural analysis

##### 2.1 Curating a dataset of high-quality experimental homomeric structures

Biological assemblies of 780 *Homo sapiens* (human), 220 *Saccharomyces cerevisiae* (yeast), and 596 *Escherichia coli* homomeric structures resolved by X-ray crystallography were obtained from the 3DComplex database (*3*, *4*). To obtain these structures, the following set of criteria was used.

(i) We selected homomers whose error rate as annotated by QSBIO was below 15% (*4*). (ii) We imposed a minimum sequence coverage of ≥ 70% for the respective UniProt sequence (*5*). (iii) Structures corresponding to artificial fusion proteins (N = 33) or proteins encoded by transposon elements (N = 12) were discarded. Finally, if more than one structure satisfied the above criteria for a particular UniProt sequence, the one with the best resolution was kept.

In addition, biological assemblies of 47 human, 15 yeast, and 15 *E. coli* homomeric proteins resolved by Electron Microscopy (EM) were added (*6*). These EM structures satisfied criteria (ii)-(iii) and were not included in the X-ray set. For *H. sapiens*, an additional set of 10 homomers, resolved by Nuclear Magnetic Resonance spectroscopy (NMR) were obtained. These NMR-homomers also satisfied criteria (ii)-(iii) and were not included in the X-ray/EM set. The final human, yeast and *E. coli* datasets comprised 837, 235, and 611 homomers.

##### 2.2 Dataset of predicted homomeric structures

We used the results from a recently developed framework that predicts homomeric structures across full proteomes (*7*). Using these predictions enabled us to map a high-quality dataset of 3584 human and 1888 *E. coli* homomeric structures to the disome enrichment data. For yeast, a total of 1414 structures were obtained.

##### 2.3 Dataset of experimental heteromeric structures

An initial set of 2440 human, 351 yeast, and 332 *E. coli* heteromeric X-ray crystallographic structures were obtained from the 3D Complex database (*3*, *4*). This dataset was combined with 1380 human, 220 yeast, and 339 *E. coli* heteromeric EM-resolved structures collected from the Protein Data Bank (*8*). These structures were further filtered to extract unique heterodimeric pairs based on the following criteria: (i) The structures covered ≥ 50% of their respective UniProt-annotated (*5*) protein lengths for both subunits. (ii) Pairs that included fusion proteins or proteins encoded by transposon elements were discarded. (iii) For a given protein pair, if more than one structure satisfied the above criteria, only the highest resolution structure was kept. The final dataset comprised 3230 human, 1728 yeast, and 628 *E. coli* unique heterodimers, encompassing 1368, 700, and 383 genes respectively.

##### 2.4 Dataset of predicted heteromeric structures

We also used a recent dataset of AlphaFold2-predicted (*9*) structures for 65,484 human binary PPI interactions (*10*). These structures were subsequently filtered in the following manner. (i) We excluded structures exhibiting steric clashes between the polypeptide chains. To that end, inter-chain $C\alpha$–$C\alpha$ distances were computed and clashes were defined as pairs of $C\alpha$ atoms closer than the sum of their van der Waals radii. Heterodimers with > 1% of $C\alpha$ atoms (summing both chains) exhibiting a steric clash were excluded. (ii) Structures with small interfaces (involving fewer than 5 residues from either chain) were also excluded. These filtering steps provided a dataset of 9349 unique heterodimeric structures covering 3465 genes. For yeast, a similar dataset of 988 heterodimeric pairs, including 1201 genes were obtained (*11*).

All the structure data for human, yeast, and *E. coli* are detailed in **Data S1, S2,** and **S3**.

**2.5 Computing structural features of protein complexes**

***Atomic contacts and protein interfaces.*** Two non-hydrogen atoms were assigned a contact if their Euclidian distance was below the sum of their van der Waals radii plus 0.5 Å (*12*). An amino acid residue exhibiting at least one intermolecular atomic contact was assigned as an interface residue. For each homomeric structure we only considered the largest interface.

***Interface residue dispersion.*** The dispersion of interface residues was measured by their average separation in the sequence, normalized by length.

$$D = \frac{1}{NL} \sum_{i=1}^{N-1} \{s(i + 1) - s(i)\} \qquad (1)$$

Where $s(i + 1)$ and $s(i)$ are the amino acid positions of the $(i + 1)$-th and the $i$-th interface residue. *N* is the total number of interface residues, and *L* is the protein length. The higher the numerical value of this parameter, the more dispersed the interface residues are relative to the protein size.

***The ratio of (inter/intra)-molecular contacts.*** For homo-/hetero-dimeric structures, we identified the intra- ($c_{intra}$) and intermolecular ($c_{inter}$) contacts. The ratio of (inter/intra) molecular contacts of interface residues were computed as described in ref. (*13*).

$$R = \frac{1}{N} \sum_{j=1}^{N} \frac{\left(c^{j}_{inter} + 1\right)}{\left(\Delta(i,j) c^{j}_{intra} + 1\right)} \qquad (2)$$

Where $j$ is the $j$-th interface residue, $c^{j}_{inter}$ is the number of intermolecular contacts for residue *j*, $c^{j}_{intra}$ is the number of intramolecular contacts for residue *j*, *N* is the total number of interface residues. The quantity $\Delta(i, j)$ represents the condition that $|s(i) - s(j)| > 4$, with $s(i)$ and $s(j)$ being the amino acid numbers in the sequence. This condition excludes intra-molecular contacts among consecutive amino acids. We add one to the denominator and numerator to avoid infinite values.

***Convex Hull analysis.*** We measured the volumes of convex hulls confining the mono- and dimeric structures. A convex hull is defined as the smallest convex geometrical enclosure confining a point-cloud. For a 3D point-cloud, such as the coordinates of the non-hydrogen atoms of a protein, the hull is a polyhedron. We employed a python implementation of Quickhull algorithm (*14*) to define polyhedron convex hulls confining monomeric and dimeric structures (**Fig. S2**). To compute the volume of space enclosed within the hull, the 3D space was split into pixels of 1 $Å^3$ size (1 Å × 1 Å × 1 Å). A point was placed at the center of each pixel. The hull volume was computed as the total number of these points that happen to be inside the hull. If the volumes of the monomeric and dimeric hull are $V_m$ and $V_d$, we computed the $V_m/V_d$ ratio.

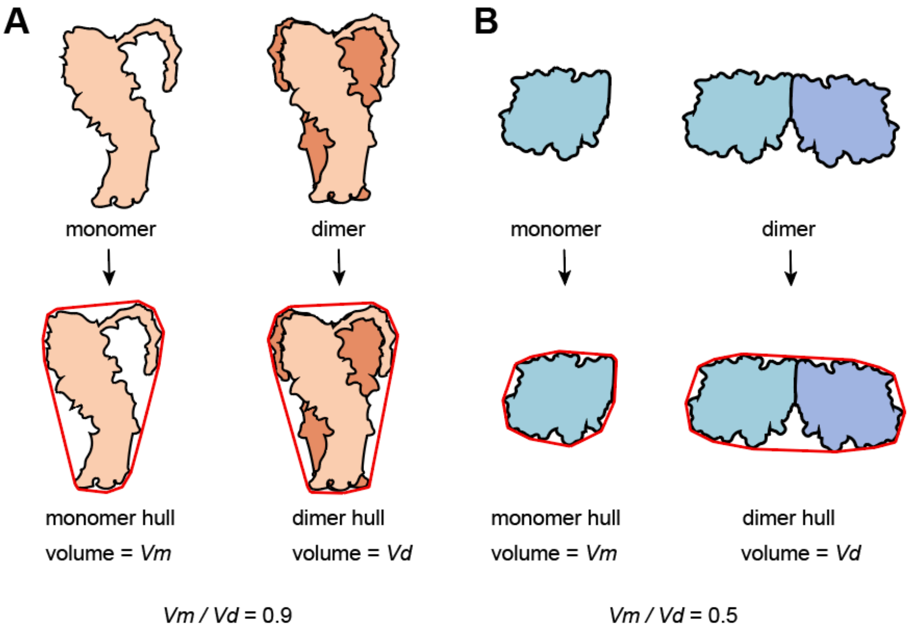


**Fig. S2**. **A schematic representation of the convex hull analysis.** A convex hull is the smallest convex polyhedron (polygon in 2D, red) confining a protein. We computed the volumes of the hulls confining the mono- and dimeric states. The fraction of the volume of the dimer hull occupied by the monomer hull is a measure of subunit intertwinement. (**A**) A homomeric complex in which the two subunits are highly intertwined, the monomer hull occupies 90% of the dimer hull volume. (**B**) The same analysis for a homomeric complex involving globular subunits interacting via a flat interface; here, the monomer hull occupies 50% of the dimer hull volume.

***Rigid Body Entanglement analysis.*** A second measure of subunit intertwinement of a dimer is the number of ways in which two subunits can dissociate as rigid bodies without causing steric clashes. To that end, for any homo-/hetero-dimer, we kept one subunit static, whereas the other one was translated by 4 Å in all directions around its center of mass. One thousand directions were uniformly sampled using the Fibonacci sphere algorithm (*15*). Thus, we iteratively translated one subunit from its center of mass to each direction and assessed steric clashes. Only backbone atoms were used to calculate clashes, which were defined as two atoms closer than the sum of their van der Waals radii. Any such shift causing >1% of all backbone atoms to clash was assigned as a forbidden dissociation direction (*FDD,* see **Fig. S3**). The fraction of *FDD*s is termed here *Rigid Body Entanglement* (*RBE*) and is a measure of subunit intertwinement. Indeed, for a highly intertwined structure, most of the 1000 attempted separation modes will be forbidden.

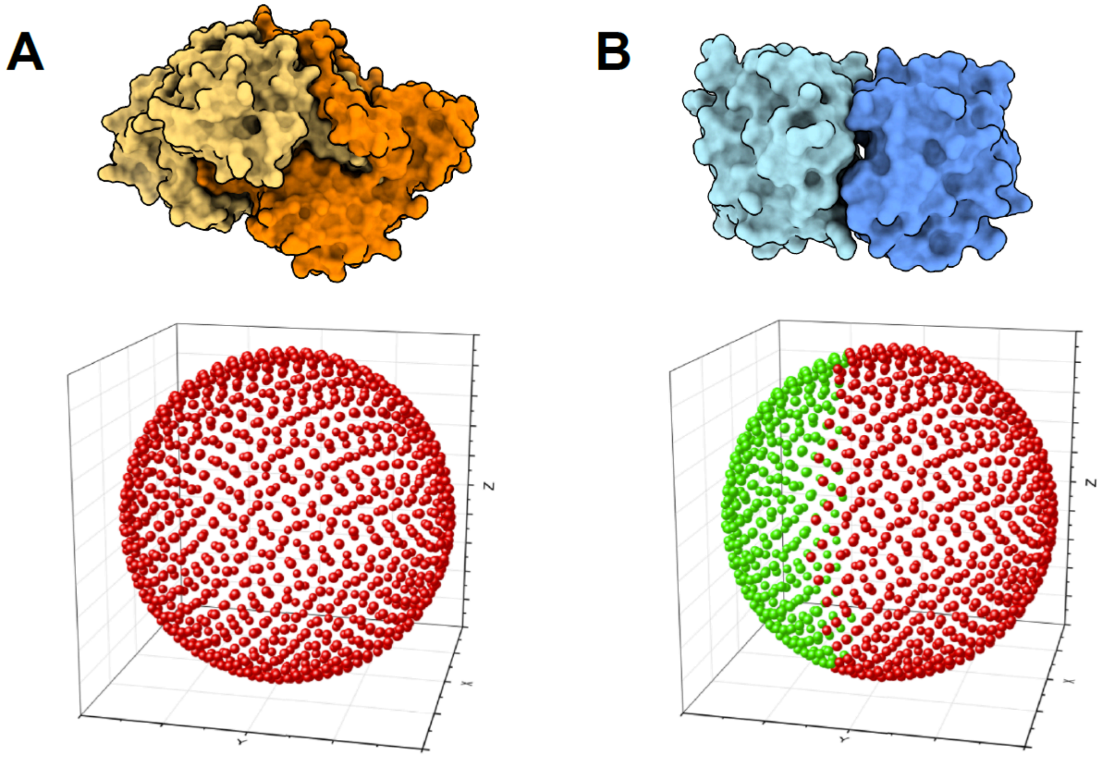


**Fig. S3**. **A schematic representation of rigid body entanglement analysis.** Here, one subunit is fixed, while the other one is shifted in all directions to test whether they can be separated as rigid bodies without causing steric clashes. The translation operation originates at the center of mass of the subunit and is performed in 1000 directions uniformly distributed on a sphere. Red points signify forbidden dissociation directions that involve significant steric clashes. Green points signify allowed dissociation directions that do not cause steric clashes. (**A**) For a homomeric complex with intertwined subunits, all dissociation modes are forbidden. (**B**) However a homomeric complex involving globular subunits is compatible with a large number of dissociation directions.

***Absolute Interface Wrapping analysis.*** As a third measure of subunit intertwinement we quantified the degree of interface non-planarity (**Fig. S4**). We fitted a plane to the 3D coordinates of the interface atoms from both subunits and calculated the atoms' sum of squared distances. This sum grows with larger and non-planar interfaces and is termed *Absolute Interface Wrapping* (*AIW*).

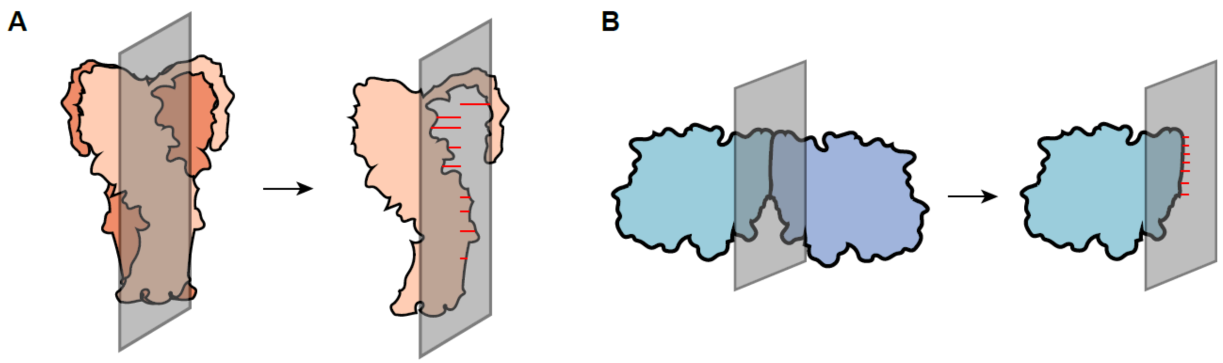


**Fig. S4. Schematic representation of the absolute interface wrapping analysis.** A 2D plane (grey) is fitted to the coordinates of interface atoms by least-squares. The root-sum-square of these distances are relatively higher for interfaces involving intertwined subunits (**A**) than for dimers involving globular subunits interacting through flat interfaces (**B**).

***Interface Size.*** We measured interface size by the number of atoms mediating inter-protein contacts.

***Measuring the non-globularity/shape of a protein.*** Solvent-exposed atoms were identified using FreeSASA (*16*) as those with ≥ 5 Å$^2$ of solvent-accessible surface area. The standard deviation of these atom's distances to the subunit center of mass served as a measure of non-globularity. For a sphere, this standard deviation would be zero.

***Measuring Absolute Contact Order.*** The absolute contact order (*ACO*) of an isolated monomer is defined as the average primary chain separation of intramolecular contacts (*17*) and is expressed as

$$ACO = \frac{1}{N}\sum_{i,j} \Delta(i,j)|s(i) - s(j)| \qquad (3)$$

Here, $N$ is the total number of intra-molecular contacts, $s(i)$ and $s(j)$ are the sequence positions of residues $i$ and $j$, and $\Delta(i,j)$ is a selection criterion that includes $i$ and $j$ in the analysis if $|s(i) - s(j)| > 4$. This criterion ensures that the contacts included in deriving *ACO* capture the topology of the protein rather than secondary structures.

***Protein Length.*** Protein length information (number of amino acids in the sequence) was derived from the UniProt sequence (*18*).

***Relationships among the structural parameters.*** We analyzed co-variation among the parameters by computing their pairwise Pearson correlations (**Fig. S5)** All parameters captured unique properties of the structures, although three parameters (*ConvexHull*, *RBE*, and *AIW),* which all capture the intertwined geometry of the complexes, co-varied the most.

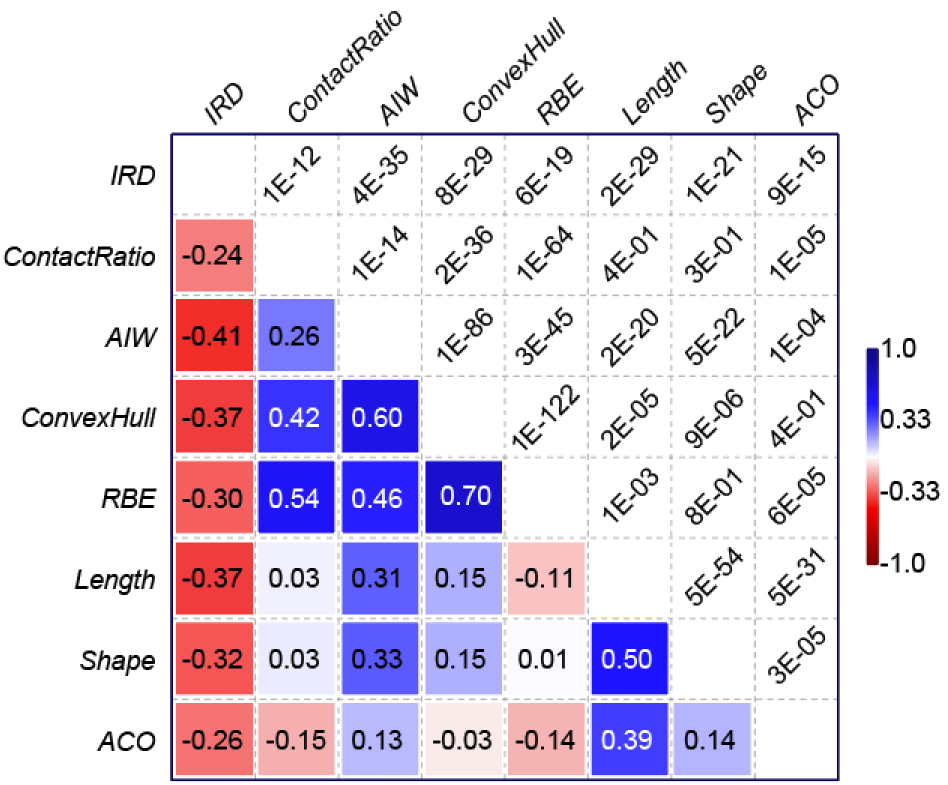


**Fig. S5**. Pearson correlations between the eight structural parameters used in this study.

#### 3. Proteomic data analysis

##### 3.1 Curating a dataset of human co-complex heteromeric pairs

Protein composition data of heteromeric human macromolecular complexes were obtained from Complex Portal (*19*) and hu.Map2 databases (*20*). The Complex Portal database currently annotates 1523 experimentally characterized heteromeric human complexes. The hu.Map2 database, on the other hand, harbors 6927 heteromeric complexes inferred from large-scale affinity purification mass spectrometry, biochemical fractionation, proximity labeling, and pulldown data. We added 3230 additional heteromeric pairs with predicted structures to this dataset, curated in *Methods 2.3*. The subunit composition data of these different sources were pulled together, resulting in a dataset of 57,769 heteromeric co-complex pairs (**Data S4**). These pairs were used for all the OMICs data analysis described below.

To assign coco and post heterodimeric pairs, these co-complex pairs were mapped to the coco and post human proteins (*Methods 1*). Pairs for which both proteins were coco were assigned as coco pairs (N = 4016), those for which both were post, were assigned as post pairs (N = 15,839).

##### 3.2 Protein melting temperature analysis

Mass spectrometry (MS) based melting temperature ($T_m$) measurements rely on measuring the abundance ratio of each protein in the soluble fraction vs pellet as a function of temperature, as unfolded proteins will aggregate and precipitate. The curve describing this relationship can serve to estimate melting temperatures, $T_m$s, (*21*) which can be compared between proteins. Alternatively, the root-mean-square deviations of the points of the curve can serve to measure similarity in melting behavior (*22*). We obtained two such datasets, the first one reports melting-temperatures of 5666 human proteins (*21*); the second reports melting-curve data for 7693 human proteins (*22*).

Using the former $T_m$ dataset, we measured a melting temperature deviation value ($\Delta T_m$) per complex composed of $N$ unique subunits as:

$$\Delta T_m = \frac{\sum_{i=1}^{N} \left| T_m^i - T_m^{mean} \right|}{N T_m^{mean}}, \qquad (4)$$

where $T_m^i$ is the melting temperature of the $i$-th subunit, and $T_m^{mean}$ is the mean melting temperature of all subunits. The smaller the $\Delta T_m$ value, the more similar the subunit $T_m$s are. We note that in this work we applied this metric to heterodimers only ($N$ = 2). Considering the second dataset of melting curves, we computed a melting curve dissimilarity score for protein pairs as the root-mean-square deviation ($RMSD_{\text{MeltCurve}}$) between the curves. Smaller values indicate similar melting behavior (*Supplementary Note 3*).

The $\Delta T_m$ and $RMSD_{\text{MeltCurve}}$ matrices (for all possible gene pairs in the human genome) are provided in **Data S5** and **Data S6**.

##### 3.3 Protein abundance data analysis

Protein abundance datasets for 13980 human proteins across 156 human cells/tissues were obtained from PaxDB (*23*). The abundance values across different datasets were merged into a matrix, where each row represents a protein and columns represent abundances across datasets. Whether two proteins are present at similar levels across cells/tissues was assessed by the Spearman correlation of their abundances (considering all the defined values for each pair).

The correlation matrix (for all possible gene pairs) is provided in **Data S7**.

##### 3.4 Protein relative degradation rate analysis

Degradation rates for human proteins have been measured by isotope labeling and measurement of the isotope "dilution" after cells are transferred to unlabelled growth medium. Dilution is quantified by MS as a function of time, from which relative degradation rates are derived (*24*). This dataset comprised relative degradation rates (σ) for 5721 human proteins. The relative degradation rate deviation (Δσ) was calculated as:

$$\Delta\sigma = \frac{\sum_{i=1}^{N} \left| \sigma^{i} - \sigma^{mean} \right|}{N\sigma^{mean}}, \qquad (5)$$

where the definitions of the different terms are the same as Equation-4. The Δσ matrix (for all possible gene pairs) is provided in **Data S8**.

#### 4. Transcriptomic data analysis

##### 4.1 Gene transcription rate analysis

Hausser et al. (*25*) analyzed high-throughput datasets of the abundance of mRNA molecules and their steady-state decay rates in the cell to infer the rates at which they are synthesized. Hence, this approach estimates mRNA production rates, which includes transcription+splicing rates ($r_{t+s}$). We collected these rates for 7793 human genes. The deviation of transcription+splicing rates ($\Delta r_{t+s}$) was measured as:

$$\Delta r_{t+s} = \frac{\sum_{i=1}^{N} \left| r_{t+s}^{i} - r_{t+s}^{mean} \right|}{N r_{t+s}^{mean}}, \qquad (6)$$

where the definitions of the different terms are the same as Equation-4. The $\Delta r_{t+s}$ matrix (for all possible gene pairs) is provided in **Data S9**.

##### 4.2 Gene expression / mRNA abundance analysis

The mRNA abundance data was collected from Bgee (*26*), which only includes data from cells/tissues not subjected to stress, disease, or genetic perturbations. In these datasets, the normalized abundance of a given mRNA (*Transcript per million*, or *TPM* score) was used. A total of 273 datasets covering 53 different human tissues were compiled and these encompass 60675 annotated human mRNAs. The mRNAs were identified by Ensembl transcript IDs and were mapped onto the human reference proteome according to UniProt mapping. In cases where multiple transcripts matched one UniProt ID, we chose the one with the highest mean abundance.

The mRNA abundance datasets were merged into a matrix and we computed pairwise Spearman correlation of the *TPM* values considering all defined values for a pair. The correlation matrix (for all possible gene pairs) is provided in **Data S10**.

##### 4.3 Translation elongation speed and initiation efficiency analysis

Lian et al. analyzed ribosomal footprinting and mRNA abundance data to derive the translation initiation efficiency (*E*) and translation elongation rates (*S*) for 8950 human genes (*27*). The deviation of translation initiation efficiency ($\Delta E$), was calculated as:

$$\Delta E = \frac{\sum_{i=1}^{N} \left| \Delta E^{i} - \Delta E^{mean} \right|}{N \Delta E^{mean}} \qquad (7).$$

The translation elongation speed dissimilarity ($\Delta S$), was calculated as:

$$\Delta S = \frac{\sum_{i=1}^{N} \left| \Delta S^{i} - \Delta S^{mean} \right|}{N \Delta S^{mean}} \qquad (8).$$

These definitions are comparable to Equation-4. The $\Delta E$ and the $\Delta S$ matrices (for all possible gene pairs) are provided as **Data S11** and **S12**.

#### 5. Prediction of co-translational assembly

##### 5.1 Developing a prediction model based on logistic-regression

To predict co-translationally assembling homomeric and hetero-dimeric pairs, we developed a logistic-regression-based probabilistic prediction model. Importantly, the identity of pairs is unambiguous for homomers, which interact with themselves, but is ambiguous for heteromers and not readily available from the experimental disome-enrichments. Therefore, we employed a three-step approach described below.

1. Infer a co-translational assembly probability from structural data for homomeric proteins with known structures.
2. Infer a co-translational assembly probability for all possible heteromeric pairs of known structures annotated as coco or post based on the experimental disome-enrichment.
3. Use steps 1 and 2 to refine the training set of heteromeric pairs, and perform a retraining that yields the final model for heteromers.

**Predicting homo-oligomers.** The training dataset included AlphaFold-modeled structures from the human homo-oligomerization Atlas (*7*) as well as structures from the PDB. All structures from the Atlas were dimers, and in cases where a homomer from PDB contained more than one interface we chose the largest one. Each homomer was annotated as coco or post based on the experimental disome-enrichment data (**Fig. S1**), which resulted in 672 coco and 1376 post homomers of known structure.

The model was trained to differentiate between these two categories based on structural features of the homomers. We employed eight features (*Methods 2.5*) and fit the dataset by logistic regression using the scikit-learn package v1.3 (*28*). The resulting model showed that eight features (interface residue dispersion (*IRD*), The ratio of (inter/intra)-molecular contacts (*ContactRatio*), Mono-/Dimer convex hull volume ratio (*CHV*), Rigid Body Entanglement (*RBE*), Absolute Interface Wrapping (*AIW*), protein non-globular shape (*Shape*), Absolute Contact Order (*ACO*), and Protein Length (*Length*)) contributed significantly to the model (*Supplementary Note 2*). We evaluated the accuracy of this model by a Receiver Operating Characteristic curve analysis using the logit probabilities as a classifier and observed an AUC of 0.95.

**First step in hetero-oligomer prediction.** We extracted all heteromeric pairs of interacting subunits from the PDB and also added model structures from ref.(*9*), as described in *Methods 2.4*. Each pair was inferred as coco when both subunits were coco in the disome enrichment data, or was inferred as post when both subunits were post in the experimental data. This resulted in 976 coco and 3796 post heteromeric pairs of known structure. We fitted a logistic regression model to this dataset using the same eight features.

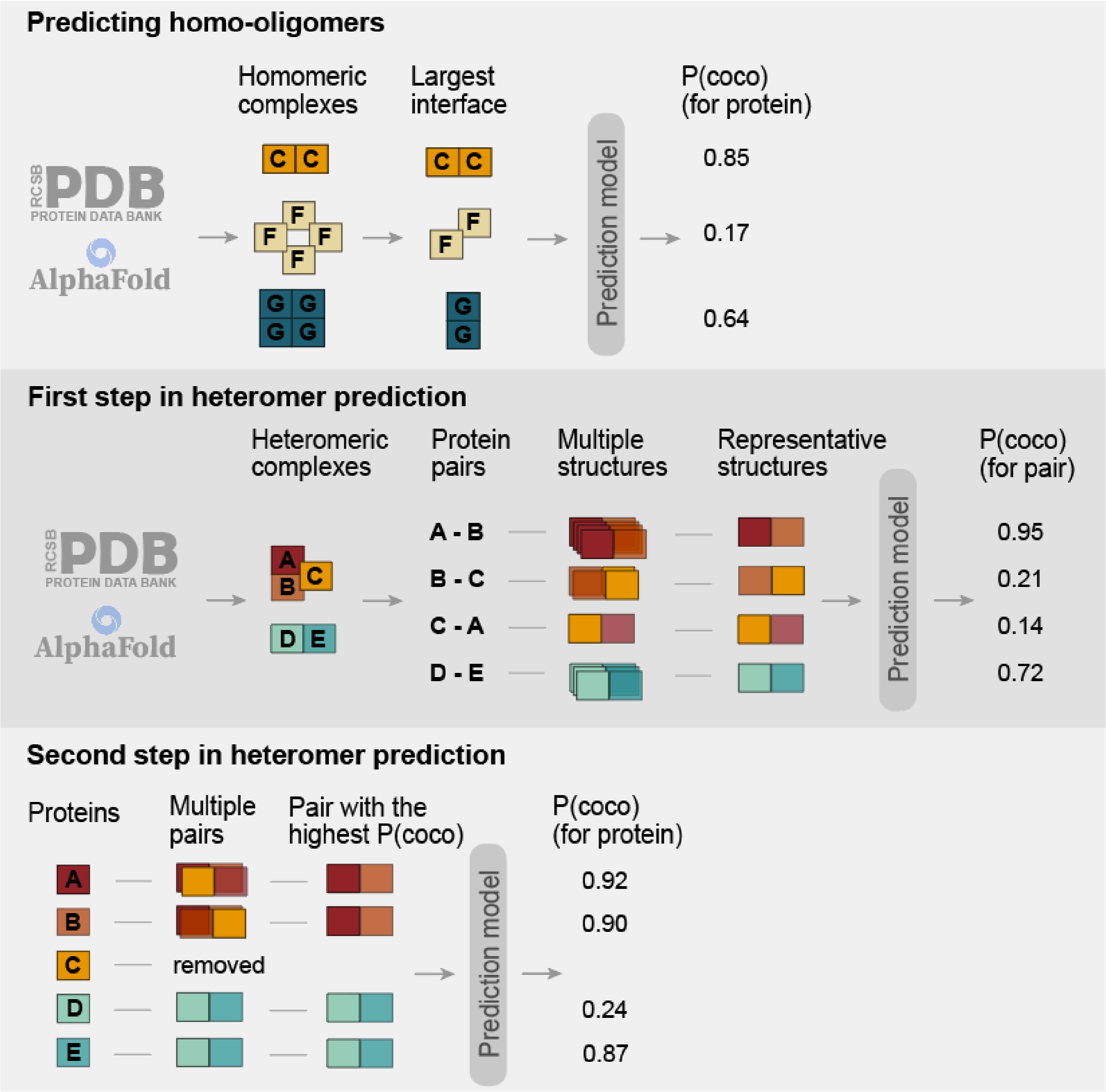


**Fig. S6**. A schematic diagram of the computational pipeline to predict co- and post-translational genes. **(A)** Inferring a co-translational assembly probability score for homomeric proteins with known structures. **(B)** Infer a co-translational assembly probability score for all possible heteromeric pairs of known structures. **(C)** Refining the training set of heteromeric pairs using homomer predictions, and performing a retraining that yields the final probability scores for heteromeric proteins.

**Second step in hetero-oligomer prediction**. Importantly, we expected the first round of assignments to include a fraction of mis-assigned pairs. For example, a complex A2B2 might appear as coco in the disome enrichment because A2 assembles co-translationally, and so does B2. In the initial set, such an AB interaction could therefore be assigned as coco, even though the heteromer formation does not involve co-translational assembly. In this second step we tried to eliminate this and other such cases by applying the following filters. (i) we removed any protein predicted as coco homomer. (ii) for each protein, we selected a single structure pair that exhibited the highest coco logit probability in the first step. Our aim with this second filter was to derive predictions for proteins and not pairs, which were more directly comparable to the disome enrichment data. These two filters resulted in 553 coco and 2113 post heteromeric proteins of known structure. Fitting this new dataset with the same eight structural features yielded the final model for heteromer co-translational assembly prediction.

We therefore derived two independent models, one applicable to homomers and the second for heteromers. Each model yields a probability for an interacting pair of known structures to assemble co-translationally. The resulting probabilities range from 0 to 1 and we considered scores ≥ 0.5 as a prediction of a positive (coco) prediction.

##### 5.2 Predicting in *E. coli*

To extend our predictions in *E. coli*, a similar three-step approach was undertaken. First, a dataset of 2384 homomeric protein structures was compiled including both PDB and AlphaFold modeled complexes (*Methods 2.1, 2.2*). In addition, a set of 628 heterodimeric pairs was compiled from PDB structures (*Methods 2.3*). The eight structural parameters computed for these structures were fed to the logistic-regression based ML models trained in human. The coco probabilities obtained in the first step were further used to filter both homomer and heteromer datasets, and also to assign a single structure for each protein, the one with the highest coco probability. A second round of predictions were made for the final, filtered structures, which were then validated by experimental disome enrichment data. These coco probabilities are available in **Data S2**.

##### 5.3 Predicting in yeast

Predictions in yeast followed the same approach as in *E. coli*. The final predictions included 2636 proteins, out of which homomeric structures were available for 1457 proteins and heteromeric structures were available for 1484 proteins. The predicted coco probabilities of these proteins are available in **Data S3**.

##### 5.4 Performance measurement

***Receiver operating characteristic curve analysis***. To evaluate the overall performance of our prediction model, we computed the receiver operating characteristic (ROC) curve. It is plotted by calculating sensitivity (True Positive Rate; TPR) against 1 − specificity (False Positive Rate; FPR) under different thresholds. The area under the ROC curve (AUC) is typically used as a metric representing the overall performance. The AUC score ranges from 0.5 to 1.0, and larger scores reflect better performance.

***Precision-Recall analysis***. The precision score of a model measures the proportion of positively predicted labels that are correct. It is particularly relevant to evaluate prediction models when the classes are imbalanced (which is the case here). It measures true positives (*TP*) relative to true positives and false positives (*FP*).

$$Precision = \frac{TP}{(FP+TP)} \qquad (9).$$

The recall measures true positives out of all positives, which include true positives and false negatives (*FN*).

$$Recall = \frac{TP}{(FN+TP)} \qquad (10).$$

##### 5.5 Infering proteome-wide numbers of co- and post-translational proteins

To obtain a genome-scale distribution of co- and post-translational proteins we combined disome enrichment data with our predictions. For the human reference proteome (*18*), the disome enrichment data annotates 2512 coco and 8547 post genes, and 3392 genes remain ambiguous (*Methods 1*). No enrichment data were available for an additional set of 5612 genes that were not detected in the experiments (no-data).

Structure-based predictions were available for a total of 6603 proteins that we classified into three categories: coco, copost, and post. Because co-translational assembly of homomers is not expected to show directionality, they were annotated as either coco or post. We then combined experimental data with these predictions to annotate human proteins. A protein was classified as coco if any evidence of it being coco existed, yielding a set of 3073 proteins.

Considering the remaining proteins, 170 were classified as copost based on structure information. These included 128 and 42 previously identified as post or having no-data (**Fig. 4A, Data S1**).

Finally, 848 and 1218 proteins previously annotated as ambiguous or having no available data were annotated as post based on their structure.

In summary, the final annotated human proteome included 3073 coco, 170 copost, 10395 post proteins, whereas 2304 and 4121 remained ambiguous or had no no-data to enable their annotation.

A similar analysis in *E. coli* resulted in 885 coco, 51 copost, and 1929 post genes, with 406 remaining ambiguous and 1132 with no-data (**Fig. 4A, Data S2**).

For yeast, no genome-scale disome enrichment data was available. Hence we used only structure-based predictions to annotate 699 coco, 153 copost, and 1784 post genes, while 3424 genes had no-data (**Fig. 4A, Data S3**).

#### 6. Selective Ribosome Profiling, SeRP

SeRP was conducted as previously described (*29*). This method enables the comprehensive profiling of interactions between factors interacting with translating ribosomes in cells. In this process, cells are flash-frozen, lyzed, and the lysate is subjected to RNase treatment to digest unprotected RNA and subsequently isolate ribosome footprints. Prior to footprint isolation, the sample is split in two, with one fraction containing all ribosome footprints (referred to as the "total translatome"), and the other undergoing affinity purification of a specific subset of ribosomes interacting with a factor of interest (referred to as the "interactome"). The protected mRNAs in both fractions are then extracted and employed for cDNA library construction, followed by deep sequencing. A comparative analysis between the total translatome and interactome samples allows for the identification of open reading frames (ORFs) preferentially associated with the enriched factor.

##### 6.1 Purification of Ribosome-Nascent-Chains (RNCs) for SeRP

RNC purification was conducted according to a well-established approach (*30*). Specifically, around 800 ml of cell culture was grown to early log phase (with an $OD_{600}$ nm of 0.5) at 30 °C in YPD medium. Cells were collected using vacuum filtration and subsequently flash-frozen. Following this, cells were lysed through cryogenic grinding using 900 µl of a lysis buffer composed of 20 mM Tris–HCl pH 8.0, 140 mM KCl, 6 mM MgCl2, 0.1 mg/ml cycloheximide (CHX), 0.1% NP-40, 1 mM PMSF, and a cocktail of protease inhibitors including Complete EDTA-free (Roche), recombinant DNaseI (Roche), leupeptin, E-64, bestatin, and aprotinin. The resulting supernatants were divided into total translatome (200 µl) and immunopurification translatome (700 µl) samples. The total translatome samples underwent RNaseI treatment (10 U per A260 nm unit) at 4 °C for 25 min, followed by layering onto sucrose cushions and subsequent ultracentrifugation at 75,000 rpm for 90 min at 4 °C. The resulting pellets were resuspended in lysis buffer. For immunopurification samples, RNaseI treatment was conducted using the same enzyme dosage in combination with approximately 50 µl of GFP-binder slurry, with an incubation period of 25 min at 4 °C.

##### 6.2 cDNA Library Preparation for Deep Sequencing

The cDNA library preparation process followed a previously reported protocol (*30*). Briefly, RNA extraction was carried out using pre-warmed acid phenol (Ambion). Subsequently, ribosomal footprints were isolated via gel electrophoresis in a 15% TBE–Urea polyacrylamide gel (Invitrogen) in 1× TBE (Ambion). After gel electrophoresis, the RNA fragments, with sizes ranging from approximately 25 to ~5 nucleotides, were stained with SYBR gold (Invitrogen) and then excised. The excised fragments underwent end repair and dephosphorylation via an incubation step with T4 polynucleotide kinase buffer and enzyme. RNA was mixed with various reagents for linker ligation and subjected to incubation at either 37 °C or 23 °C. Following reverse transcription, hydrolysis of RNA was carried out using NaOH, and the resulting products were circularized by treatment with CircLigase enzyme. The circularized DNA was utilized for PCR amplification, and the amplified products were subjected to quality control before being sequenced on a HiSeq 2000 (Illumina) platform.

##### 6.3 Data Analysis

Processing of sequenced reads involved established tools such as Cutadapt, Bowtie2, and Tophat2, as well as customized Python scripts tailored for *S. cerevisiae*, as previously described (*30*). Analyses were based on at least two biologically independent replicates.

#### 7. RNA Immunoprecipitation qPCR

Our RIP-qPCR protocol is an adapted version of previously published methods (*31–33*) tailored for robust RNA immunoprecipitation coupled with quantitative polymerase chain reaction (RIP-qPCR) experiments. The procedure entails the capture and quantification of RNA molecules associated with specific protein targets.

##### 7.1 Cell Culture and Lysis

Overnight cultures were cultivated in a YPD medium. These cultures served as the basis for inoculating 100 mL of fresh YPD to achieve an initial $OD_{600}$ of 0.035. Expression cultures were then grown at 30 °C and shaked at 200 rpm until reaching an $OD_{600}$ of 0.5–0.6. Cells were then harvested via 30 sec centrifugation at 3000 × g followed by resuspension in 0.5 mL YPD for a 15 min recovery before being flash-frozen in liquid nitrogen.

##### 7.2 Cell Lysis and Bead Binding

Frozen sampled were supplemented with 0.5 mL of frozen high-salt lysis buffer (comprising 20 mM Tris-HCl, pH 8.0, pH 7.5, 140 mM KCl, 10 mM $MgCl_2$, 1 mM PMSF, 0.1 % NP-40, cOMPLETE EDTA-free protease inhibitor (Roche), 0.02 U/uL DNaseI and either 0.1 mg/mL CHX (Sigma-Aldrich) or 40 mM EDTA (Sigma-Aldrich)). Under cryogenic conditions, cells were grinded by CryoMilling (Retsch) at 30 Hz for 2 min. The lysate was thawed, transferred into 1.5 mL tubes, and cleared at 20,000 × g at 4 °C for 3 min. The cleared supernatant was then added to equilibrated Protein A resin conjugated to antibodies against HA (rabbit), along with 0.1 U/µL Ribolock (Invitrogen) to inhibit RNA decay. The lysate was incubated with the beads at 4 °C for 1 hr with end-to-end mixing.

##### 7.3 Bead Washing and RNA Extraction

Following incubation, the beads were washed via sequential centrifugation at 500 g and 4 °C for 5 min. Beads were washed three times with 1 mL of wash buffer A (20 mM Tris-HCl, pH 8.0, 140 mM KCl, 20 mM MgCl2, 0.1 % NP-40, cOMPLETE EDTA-free protease inhibitor, and 0.1 mg/mL CHX) for 1 min each by end-to-end mixing. This was followed by two washes with wash buffer B (20 mM Tris-HCl, pH 8.0, 500 mM KCl, 20 mM MgCl2, 0.01 % NP-40, cOMPLETE EDTA-free protease inhibitor, and 0.1 mg/mL CHX) for 1 min and 4 min, respectively. The beads were finally resuspended in 500 µL of 10 mM Tris-HCl, pH 8.0.

##### 7.4 RNA Extraction and Precipitation

RNA extraction was initiated by adding 40 µL of 20% SDS and 750 µL of pre-warmed phenol-chloroform-isoamyl alcohol (PCI, 65 °C, Invitrogen). The mixture was incubated at 65 °C, 1,400 rpm for 5 min, followed by rapid cooling on ice for 10 min. After centrifugation at 15,000 g for 10 min, the aqueous phase was subjected to a second round of PCI extraction at room temperature for 5 min. A diethyl ether wash eliminated residual PCI, and the remaining solvent was evaporated in a Speedvac (Eppendorf).

##### 7.5 RNA Precipitation and cDNA Synthesis

RNA precipitation was achieved by adding 3 M NaOAc, pH 5.5, to attain a final concentration of 0.3 M. 2.5 µL of Glycoblue (Invitrogen) and an equal volume of isopropanol were added to the precipitate. The mixture was then placed in a −80 °C freezer overnight. Subsequent centrifugation at 15,000 g and 4 °C for 90 min yielded a pellet, which was washed with 70% EtOH, dried in a Speedvac (Eppendorf), and resuspended in 20 µL of 10 mM Tris-HCl, pH 8.0. RNA precipitations generally yielded 150–250 ng/µL of RNA.

**7.6 cDNA Synthesis and qPCR**

For cDNA synthesis, 500 ng of RNA was used, and cDNA was synthesized following the manufacturer's PrimeScript™ RT reagent kit (Takara) instructions, including an optional gDNA eraser.

Real-time qPCR was conducted using the PerfeCTa SYBR ® Green FastMix (Quantabio) according to the manufacturer's protocol. The qPCR was performed using the QuantStudio 1 cycler (Applied Biosystems, 95 °C: 30 sec; 40 cycles: 95 °C: 3 sec, 60 °C: 30 sec). Images were taken every cycle within the annealing/extension step. All qPCR assays were performed in technical triplicates, and each experiment was analyzed using the QuantStudio analysis software (v1.5.1). Briefly, quality assessment was performed within the QuantStudio software and if suggested by the software individual technical replicates were omitted. Experiments in which two technical replicates were omitted did not pass our quality control filter.

Ct-values were calculated by 2nd derivation by the QuantStudio software (ThermoFisher). To calculate relative gene expression ΔCt-values were calculated by the following formula:

$$Gene\ expression\ =\ 2^{\Delta C_T}, \Delta C_T = C_T(target)\ -\ C_T(Housekeeping) \qquad (11).$$

$$Enrichment\ =\ 2^{\Delta\Delta C_T},\ \Delta\Delta C_T =\ \Delta C_T(CHX\ treated\ lysate)\ -\ \Delta C_T(EDTA\ treated\ lysate) \qquad (12).$$

For normalization Act1 mRNA was used as a housekeeping gene. Samples were analyzed in triplicates for each primer pair as technical replicates. For calculation of gene expression the average of those three replicates was used. For primer sequences see **Table S1**. Amplicons were designed with a size of 150-200 nt.

**Table S1**. The primer sequences used for RIP-qPCR experiments.

| No. | Target Gene | Primer Name | Sequence |
|---|---|---|---|
| AS342 | RPB1 | RPB1 qPCR Forward | ACTGGATCGTGCAGCAATGA |
| AS343 | RPB1 | RPB1 qPCR Reverse | GACGAACAACACGACAACGG |
| AS346 | RPB2 | RPB2 qPCR Forward | TGCCATTTACCGCAGAAGGT |
| AS347 | RPB2 | RPB2 qPCR Reverse | AGTGCCGCGACTTTACTCAA |
| AS350 | RRN6 | RRN6 qPCR Forward | ACTGCGATACGCCAGTTTCA |
| AS351 | RRN6 | RRN6 qPCR Reverse | GCATCGGCCTCGAAGAGTTA |
| AS354 | RRN7 | RRN7 qPCR Forward | GCGCACAACCACAAGTGAAT |
| AS355 | RRN7 | RRN7 qPCR Reverse | CTCCTCCTCAGACCACTGGA |
| AS358 | SKI2 | SKI2 qPCR Forward | CGTGGTGTTGTCTGGGAAGA |
| AS359 | SKI2 | SKI2 qPCR Reverse | GTGTTAGGAACGGTGGCAGA |
| AS362 | SKI3 | SKI3 qPCR Forward | CCTGGGGGTACATTTGCCAT |
| AS363 | SKI3 | SKI3 qPCR Reverse | GCAGCATTGTGGTCCTTTGG |

#### 8. Single-molecule fluorescence in situ hybridization

##### 8.1. Principle

Single-molecule fluorescence in situ hybridization (smFISH) consists of a large number (20 to 40) fluorescently labeled probes complementary to a target RNA. These probes can be visualized by microscopy (*34*). By designing multiple sets of probes of different colors, multiple targets can be simultaneously tracked. We hypothesized that co-translational assembly should involve mRNA co-localization, which we probed by smFISH.

##### 8.2. Probe Design

We used Stellaris FISH probes (*35*), designed using the online tool (URL: biosearchtech.com/support/tools/design-software/stellaris-probe-designer, accessed on 13/08/2023) according to manufacturer recommendations. Six predicted co-translational pairs were selected for experiments: Rpb1/2, Ski2/3, Ret1/Rpo31, Gcd10/14, Yku70/80, and Nuf2/Ndc80. Since specific target binding is critical to our experiments, we analyzed the melting temperature ($T_m$) of each probe and the abundance of potential off-targets. Transcript abundance values were collected from Expression Atlas (*36*). Off-target binding was examined by sequence similarity (nBLAST (*37*)) against *S. cerevisiae* transcriptome (*38*). For each BLAST-hit, binding $T_m$s were predicted using the Bio.SeqUtils.MeltingTemp module of BioPython (*39*). We excluded probes with apparent off-target sites in mRNAs with an abundance similar or higher than that of the target. For each pair, two distinct fluorophores were used, CAL Fluor Red 590 and Quasar 670. In addition, we designed two positive controls: the 5′- and 3′- halves of Rpb1 and Rpo31 mRNAs. Two probe sets with different fluorophores were designed to target two halves of these mRNAs. The final set of probes (29-40 probes per target) are detailed in **Data S13**.

##### 8.3. smFISH experiments

For RNA-hybridization, we used the protocol from Nair et al., (*40*) with some modifications. BY4741 WT yeasts were grown to the early-log phase ($OD_{600}$ = 0.3-0.4) and then incubated with cycloheximide (100 µg/ml final concentration; Sigma-Aldrich) for 15 minutes. Cells were then fixed with formaldehyde (4% final concentration), and incubated at RT for 45 min and which they were pelleted (1600 g, 4 min, 4°C) and washed twice with ice-cold Buffer B (0.1 M potassium phosphate buffer, pH 7.5, containing 1.2 M sorbitol; Sigma-Aldrich). We then digested the cell wall in 1 ml of freshly prepared spheroplast buffer (Buffer B supplemented with 20 mM ribonucleoside vanadyl complexes (Sigma-Aldrich), 20 mM β-mercaptoethanol (Sigma-Aldrich), and lyticase (A&A Biotechnology) 30 U per 50 ml of cells) for 35 min at 30°C. The spheroplasts were centrifuged for 5 min at a low speed (500 g) at 4°C, and washed twice with ice-cold Buffer B. Finally, the cells were incubated with 70% ethanol overnight at −20°C. The next day, cells were washed twice with 2x SSC buffer (0.3 M sodium chloride, 30 mM sodium citrate; Fisher BioReagents), followed by incubating with wash buffer (2x SSC with 10% formamide; Sigma-Aldrich), for 15 min at room temperature. Then, 100 µl of hybridization buffer (2x SSC, 10% dextran sulfate, 10% formamide, 2 mM ribonucleoside vanadyl complexes, 200 µg/ml *E. coli* tRNA, and 40 µg/ml bovine serum albumin; Sigma-Aldrich) containing CAL Fluo Red 590 and Quasar 670 probe sets, 125 nM each. The samples were incubated overnight at 37°C. The next day, samples were washed twice with preheated wash buffer at 37°C for 15 min. And washed once with 1 ml 2x SSC 0.1% Triton X-100 at room temperature for 5 min. Then the samples were washed twice with 1x SSC, stained by DAPI, and resuspended in 1x SSC.

**8.4. Imaging**

For imaging, samples were diluted 10 times in a 1x SSC buffer and transferred to a 96-well optical plate (30 µl of final volume). Cells were imaged with an Olympus IX83 microscope coupled to a Yokogawa CSU-W1 spinning-disc confocal scanner and dual prime-BSI sCMOS cameras (Photometrix). The 16-bit images were acquired with brightfield and three fluorescence channels of 500 ms exposure each: one for CAL Fluor Red 590 (561 nm laser; 609-54 nm filter, Chroma ET/m), one for Quasar 670 (640 nm laser; 700-75 nm filter, Chroma ET/m), and one for DAPI (405 nm laser; 700-75 nm filter, Chroma ET/m). One multiband dichroic mirror was used for the three illuminations. Imaging was performed with a 60x/1.42 numerical aperture (NA), oil-immersion objective (UPLSAPO60XO, Olympus) and with a 100x/1.45 numerical aperture (NA), oil-immersion objective (UPLXAPO100XO, Olympus). Automated imaging was performed with a motorized XY stage onto which a piezo-stage (Mad City Labs) is mounted. For each sample, 20 fields were acquired.

**8.5. Image analysis**

Cells were identified and segmented using a custom approach (*41*) in ImageJ/FIJI (*42*). An overlay of CAL Fluor Red 590 and Quasar 670 channels revealed co-localization spots/punctae in the cell. The extent of co-localization, per field of view, was quantified as the number of overlapping punctae within identified cells, normalized by the total number of punctae in the same cells. The number of overlapping punctae was obtained by counting the number of regions (minimum size of 5 pixels) after multiplying the two binary images of both channels' punctae. The mean of this ratio, over all 20 fields, was obtained for each pair and the positive controls.

#### 9. Genetic perturbation experiments

##### 9.1. Plasmids and strains

For overexpression and deletions of protein pairs, strains from the previously described C-SWAT library (*43*) with C-terminally mNeonGreen-tagged proteins were made heat shock competent (*44*). For overexpression experiments, these C-SWAT strains were transformed with 250 ng p14x plasmids (*45*) encoding the second protein with a C-terminal mScarlet-I (*46*) tag as well as a G418 selection cassette. These plasmids sequences were designed and ordered from Twist Bioscience. For deletions, the G418 selection cassette was PCR amplified using PrimeSTAR Max DNA Polymerase (Takara Bio) with custom primers (Sigma) that included a 100 bp long homology to the genome immediately before/after the start-/stop codon of the target gene. The PCR product was treated with DPN1 (NEB), purified using AMPure XP beads (Beckman Coulter), and transformed into the respective competent C- SWAT strains. The correct insertion of the G418 cassette and the successful knockout were verified via PCR (Bio-ReadyMix, Biolab-Biology Ltd.) with primers aligning up- and downstream of the cassette. All plasmids and strains generated in this study are listed in **Data S14 and S15**, respectively.

##### 9.2. Imaging

For imaging the overexpression and knockout strains, cells were grown for 3 days to saturation. Subsequently, 1 μL of the culture was transferred to 35 μL fresh media (SD media supplemented with nourseothricin and/or G418), and grown 6 h at 30°C prior to imaging. Cells were imaged, segmented, and the abundance of visible foci were analyzed as described elsewhere (*47*).
