## Supplementary Notes for "Structural determinants of co-translational protein complex assembly"

### The structure of protein complexes underlies co-translational assembly

**This Supplementary file includes:**

Supplementary Notes 1-8
Supplementary Figures S7 to S19
Supplementary Tables S2 to S5
Supplementary References

**Supplementary Datasets for this manuscript include the following:**

**Data S1**. Structure Dataset for *Homo sapiens*
**Data S2**. Structure Dataset for *Escherichia coli*
**Data S3**. Structure Dataset for *Saccharomyces cerevisiae*
**Data S4**. Human co-complex subunit pairs
**Data S5**. Delta-Melting-Temperature matrix for human
**Data S6**. Melting-Curve-Dissimilarity matrix for human
**Data S7**. Protein-Abundance-correlation matrix for human
**Data S8**. Delta-Protein-Degradation-Rate matrix for human
**Data S9**. Delta-Transcription-Rate matrix for human
**Data S10**. Gene-Expression-correlation matrix for human
**Data S11**. Delta-Translation-Initiation-Efficiency matrix for human
**Data S12**. Delta-Translation-Elongation-Speed matrix for human
**Data S13**. Fluorescent probe sequences used for smFISH experiments
**Data S14**. Plasmids generated for genetic perturbation experiments
**Data S15**. Yeast strains generated for genetic perturbation experiments

Supplementary data files are available in FigShare (DOI: 10.6084/m9.figshare.24311917).

### Supplementary Notes

#### Note 1. Topological features of human coco and post proteins

##### 1.1 Topological features of human homomers with experimental structures

As discussed in **Fig. 1**, topology underlies co- and post-translational modes of complex assembly. The former typically includes subunits intertwined in 3D space, whereas the latter includes globular subunits interacting via small, flat interfaces. We computed additional features to those shown in **Fig. 1** (*Supplementary Methods 2*) and we depict their distribution among coco and post homomer subunits in **Fig. S7**.

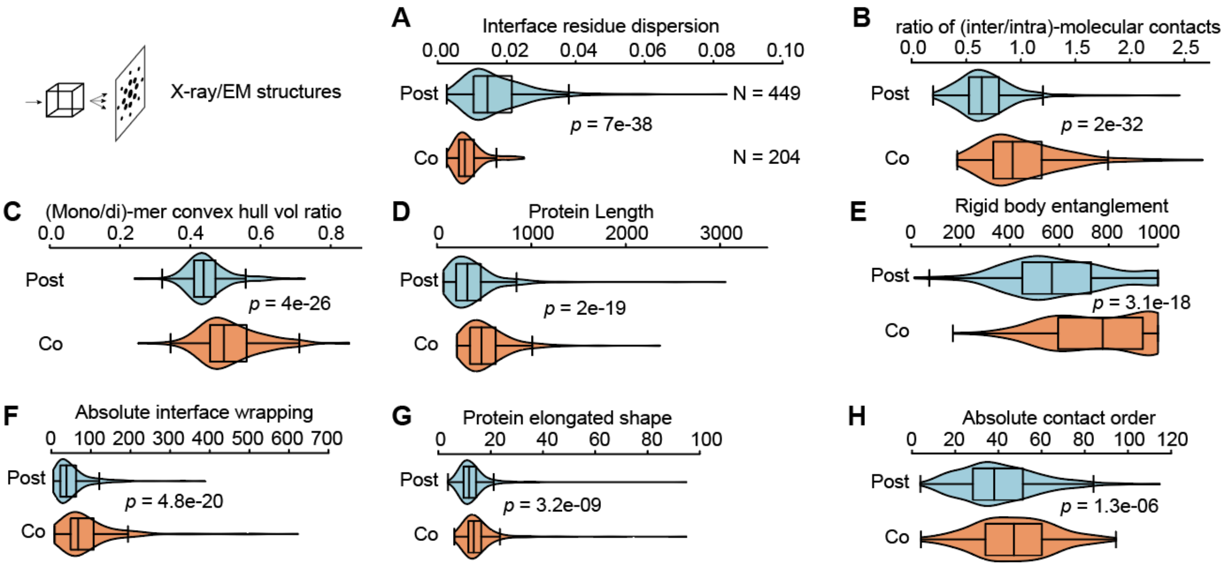


**Fig. S7. Coco and and post subunits from homomeric structures show differences in several topological features.** Panels (**A**)-(**D**) are identical to **Fig. 1**(**B**)-(**E**). Plot features follow Fig 1B. (**E**) Violin plots comparing the rigid body entanglement of co- (orange) and post-translational (cyan) homomers with available experimental structures. (**F**) Same as (A), but for absolute interface wrapping. (**G**) Same as (A), but for non-globularity of monomer shapes. (**H**) Same as (A), but for absolute contact order of monomers.

#### 1.2 Topological features of human heteromers with experimental structures

We examined whether the topological differences seen in homomers generalize to coco and post heteromeric pairs. A dataset of human heterodimeric pairs with experimental 3D structures (*Supplementary Methods 2*) was mapped to human co- and post-translational genes (*Supplementary Methods 1*). As plotted in **Fig. S8**, the topological differences between co- and post-translational heteromeric pairs depicted similar trends as homomers. This analysis includes an obvious pitfall that two co-translational proteins being in 3D contact does not necessarily mean they assemble on each-other.

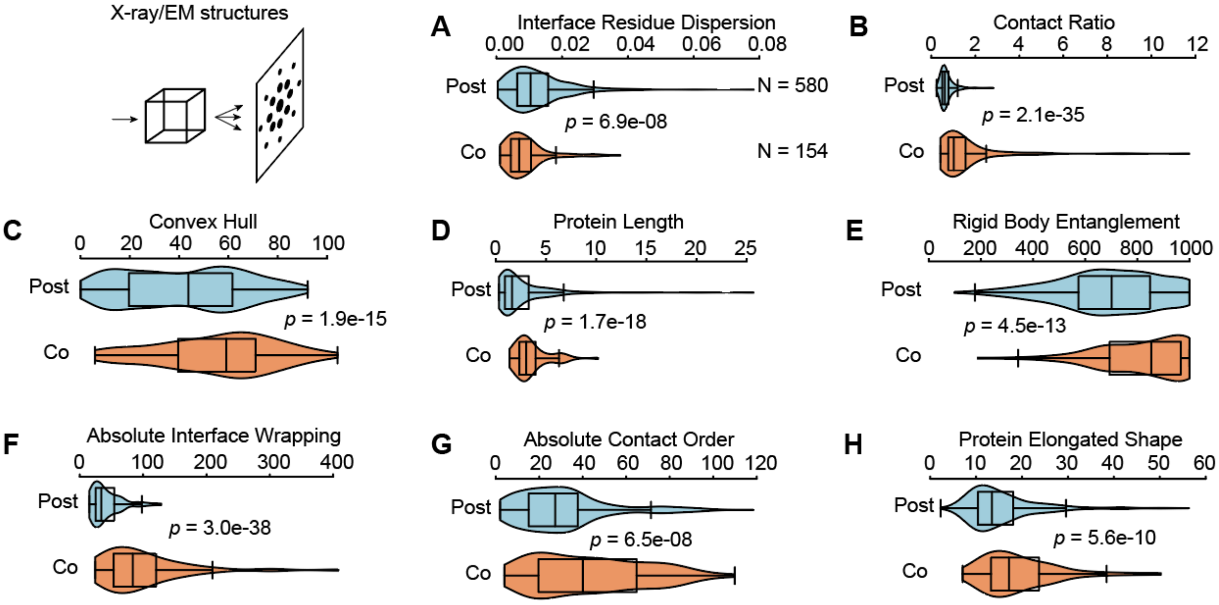


**Fig. S8. Coco and- and post subunits from heteromeric structures show differences in several topological features.**
(**A**) Violin plots comparing the interface residue dispersion of coco (orange) and post (cyan) hetero-dimers with available experimental structures. Plot features follow Fig 1B. (**B**) Same as (A) for the ratios of inter- and intra-molecular atomic contacts of the interface residues. (**C**) Same as (A) for the percentage of volume of the dimer convex hull that is occupied by that of the monomer. (**D**) Same as (A) for the protein sequence length. (**E**) Same as (A) for rigid body entanglement. (**F**) Same as (A) for absolute interface wrapping. (**G**) Same as (A) for absolute contact order of the monomers. (**H**) Same as (A) for non-globularity of monomer shapes.

#### 1.3 Topological features of human homomers with predicted structures

We examined the topological differences between co- and post- homomers (**Fig. 1** and **S7**) on larger genome-scale using predicted complexes' structures (*Supplementary Methods 2*). **Fig. S9** highlights that topological differences were further amplified in these predicted structures.

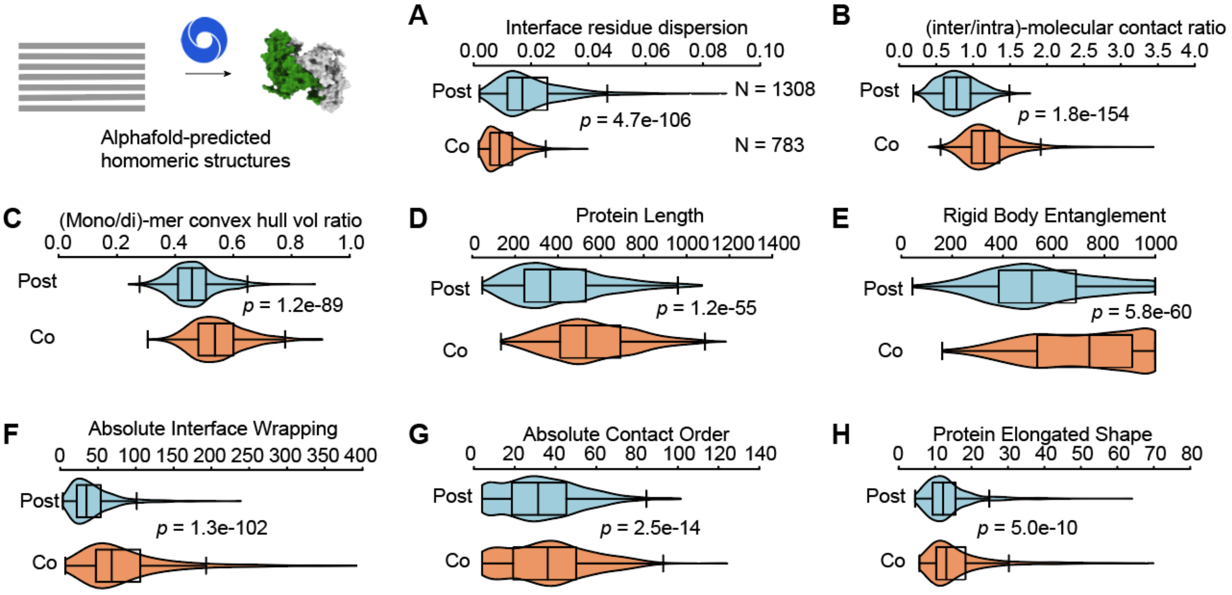


**Fig. S9**. **Coco and post subunits from homomeric structures show differences in several topological features.** (**A**) Violin plots comparing the interface residue dispersion of co- (orange) and post-translational (cyan) homomers with available experimental structures. Plot features follow Fig 1B. (**B**) Same as (A) for the ratios of inter- and intra-molecular atomic contacts of the interface residues. (**C**) Same as (A) for the percentage of volume of the dimer convex hull that is occupied by that of the monomer. (**D**) Same as (A) for the protein lengths (#amino acids). (**E**) Same as (A) for rigid body entanglement. (**F**) Same as (A) for absolute interface wrapping. (**G**) Same as (A) for absolute contact order of the monomers. (**H**) Same as (A) for non-globularity of monomer shapes.

##### 1.4 Topological features of human heteromers with predicted structures

We examined the topological differences between co- and post- heteromeric pairs (**Fig. S8**) on a genome-scale, using predicted complexes' structures (*Supplementary Methods 2*). **Fig. S10** highlights that topological differences were retained in these predicted structures.

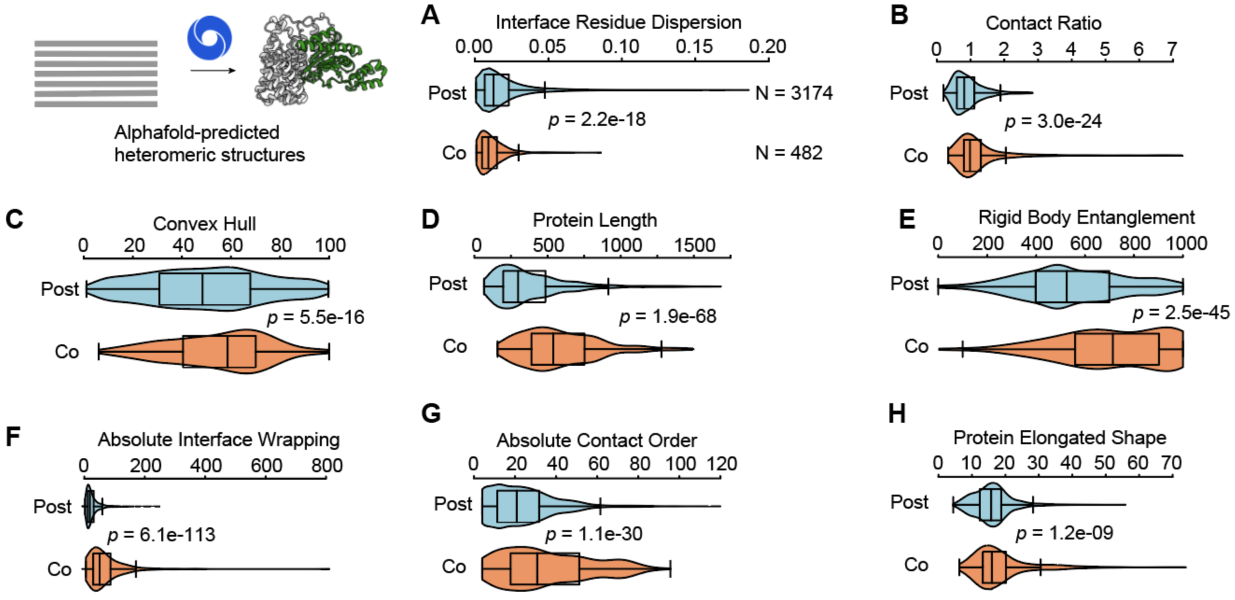


**Fig. S10**. **Coco and- and post subunits from heteromeric structures show differences in several topological features.**
(**A**) Violin plots comparing the interface residue dispersion of co- (orange) and post-translational (cyan) hetero-dimers with predicted structures. Plot features follow Fig 1B. (**B**) Same as (A) for the ratios of inter- and intra-molecular atomic contacts of the interface residues. (**C**) Same as (A) for the percentage of volume of the dimer convex hull that is occupied by that of the monomer. (**D**) Same as (A) for the protein lengths (#amino acids). (**E**) Same as (A) for rigid body entanglement. (**F**) Same as (A) for absolute interface wrapping. (**G**) Same as (A) for absolute contact order of the monomers. (**H**) Same as (A) for non-globularity of monomer shapes.

#### Note 2. Assessing the performance of the prediction model in human

##### 2.1 Parameter choice

First we developed an Ordinary Least Square (OLS) Regression model using nine parameters. These parameters included interface residue dispersion (*IRD*), The ratio of (inter/intra)-molecular contacts (*ContactRatio*), Mono-/Dimer convex hull volume ratio (*CHV*), Rigid Body Entanglement (*RBE*), Absolute Interface Wrapping (*AIW*), Interface size, protein non-globular shape (*Shape*), Absolute Contact Order (*ACO*), and Protein Length (*Length*). Because the nine structural parameters are not mutually independent, two-tailed t-tests were carried out to measure their contributions to the regression using the Python package statsmodel version 0.14.0 (*1*). As enlisted in **Table S2**, except interface size, the remaining eight parameters contributed significantly to the regression ($p < 0.05$).

**Table S2**. The *p*-values of the t-tests assessing which parameters significantly contribute to the regression model.

| Training Parameters | p-value |
|---|---|
| Shape | 0.034 |
| Absolute Interface Wrapping*, AIW* | 0.001 |
| Mono/Di-mer Convex Hull Volume ratio, *ConvexHull* | 0.000 |
| Rigid Body Entanglement, *RBE* | 0.000 |
| Interface Size | 0.646 |
| inter/intra-molecular contact ratio, *ContactRatio* | 0.000 |
| Interface residue dispersion, *IRD* | 0.000 |
| Absolute contact order, *ACO* | 0.000 |
| Protein Length | 0.002 |

To test whether interface size indeed increases the prediction accuracy in the absence of any size-related parameter, we compared the predictive power of two models, one trained on three parameters *RBE*, *IRD*, and ContactRatio (**Fig. S11A**), and the other one trained on *RBE*, *IRD*, ContactRatio, and InterfaceSize (**Fig. S11B**). The latter depicted a minor increment in prediction accuracy (AUC increased from 0.920 to 0.923), reiterating the fact that InterfaceSize did not contribute significantly to the regression model.

Large interface areas were previously shown to associate with co-translational assembly (30). The above analyses indicate that the underlying determinant of that association is subunit interaction geometry, such that intertwined subunits of co-translational complexes exhibit larger interface areas. Based on these results, we developed a logistic-regression based prediction model based on the remaining eight parameters, excluding the InterfaceSize.

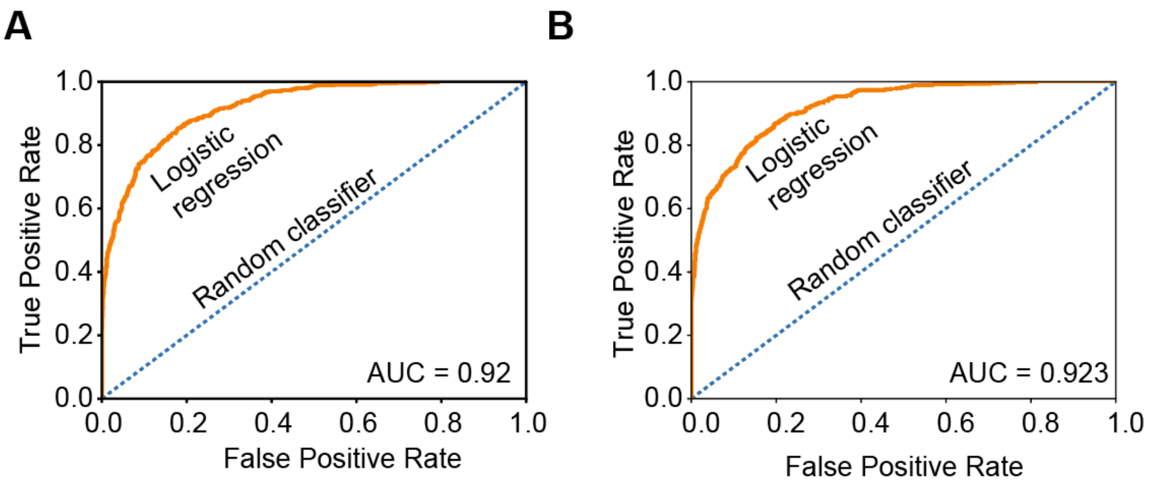


**Fig. S11**. The receiver operating characteristic (ROC) curve for predicting co- and post-translational modes of homomer assembly, for a logistic regression-based prediction model trained on **(A)** *RBE*, *IRD*, and ContactRatio and **(B)** *RBE*, *IRD*, ContactRatio, and InterfaceSize.

#### 2.2 Performance of the prediction model in human

The prediction accuracy of our logistic-regression based machine learning model was examined by two measures. The Area under the curve or AUC of the Receiver Operating Characteristic curve (**Fig. 3B**), and that of Precision-Recall curve (**Fig. S12**).

The contributions of the individual topological parameters are enlisted in **Table S3**.

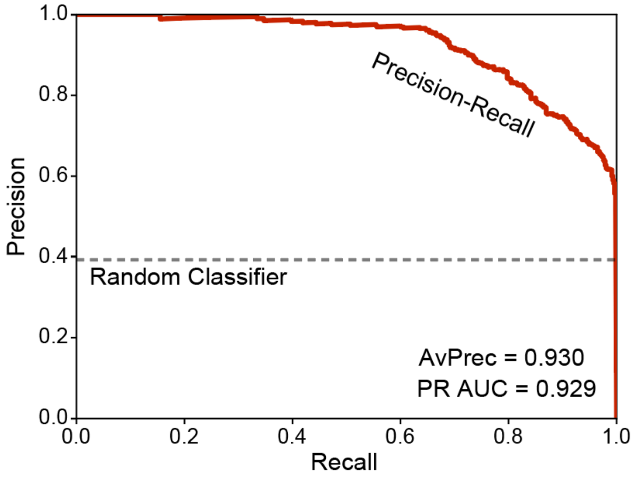


**Fig. S12**. Precision-Recall curve for predicting co- and post-translational homomers. The Precision-Recall AUC and Average Precision are mentioned.

**Table S3**. The ROC-curve AUC, and the Average Precision for individual parameters for predicting co- and post-translational homomers.

| Training Parameters | *AUC* | *AvPrec* |
|---|---|---|
| Shape, *AIW*, *ConvexHull*, *RBE*, *CntctRatio*, *IRD*, *ACO*, Length | 0.951 | 0.930 |
| Shape | 0.608 | 0.435 |
| Absolute Interface Wrapping, *AIW* | 0.793 | 0.645 |
| Mono/Di-mer Convex Hull Volume ratio, *ConvexHull* | 0.772 | 0.606 |
| Rigid Body Entanglement, *RBE* | 0.733 | 0.590 |
| inter/intra-molecular contact ratio, *CntctRatio* | 0.843 | 0.727 |
| Interface residue dispersion, *IRD* | 0.812 | 0.650 |
| Absolute contact order, *ACO* | 0.564 | 0.386 |
| Protein Length | 0.711 | 0.490 |

#### 2.3. Performance of the prediction model in *Escherichia coli*

The accuracy of predictions in *E. coli* was examined as described above.

**Fig. 3D** highlights the accuracies for homomers: AUC = 0.911, Av. Precision = 0.787.

**Fig. S14** highlights the accuracies for heteromers: AUC = 0.841, Av. Precision = 0.644.

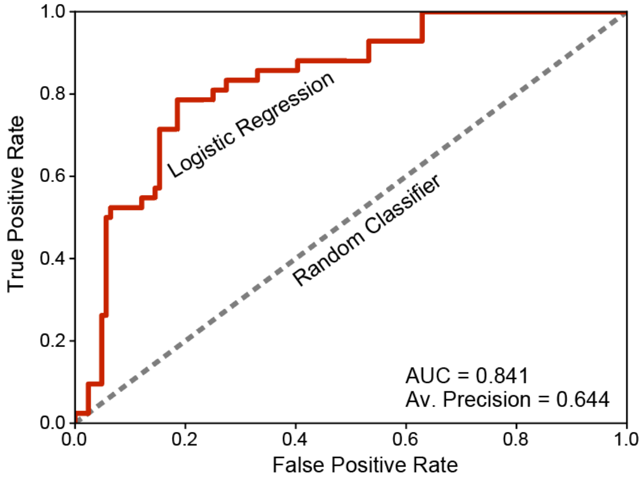


**Fig. S13**. ROC curve for predicting co- and post-translational heteromeric proteins in *E. coli*.

#### Note 3. Melting Curve Similarity of coco and post heteromeric pairs

We analyzed a proteome-scale dataset of protein melting temperatures (*2*) and found that co-translational heteromeric pairs exhibit more similar $T_m$s than post-translational pairs (**Fig. 1**). Here we reproduced these results using an independently generated dataset of protein melting curves (*3*) (*Supplementary Methods 3*, **Fig. S14**).

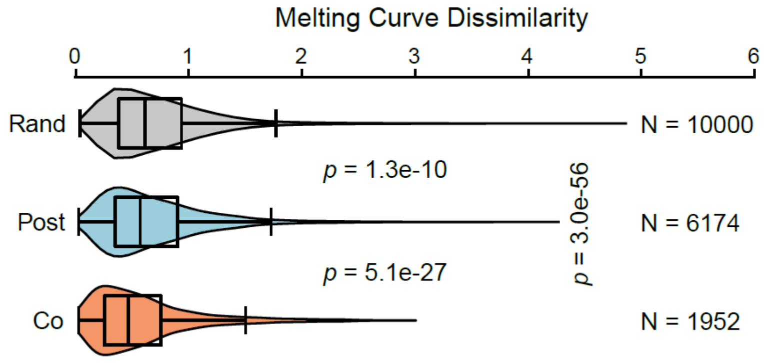


**Fig. S14**. Violin plots depicting the root-mean-square deviation of protein melting curves for coco (orange), post (cyan) and random protein pairs (gray).

#### Note 4. OMICs data analysis and directionality of co-translational assembly

As depicted in Fig. 2, the rates of various core cellular processes are relatively more synchronized for co-translational heteromeric pairs than for post. Here, we show that those differences are not only retained but further amplified once structural information and directionality of assembly is invoked. Here in **Fig. S15**, the rates of core cellular processes are compared for human heteromeric pairs with 3D structures, classified into three groups: coco (bidirectional co-translational), copost (unidirectional co-translational), and postpost (post-translational).

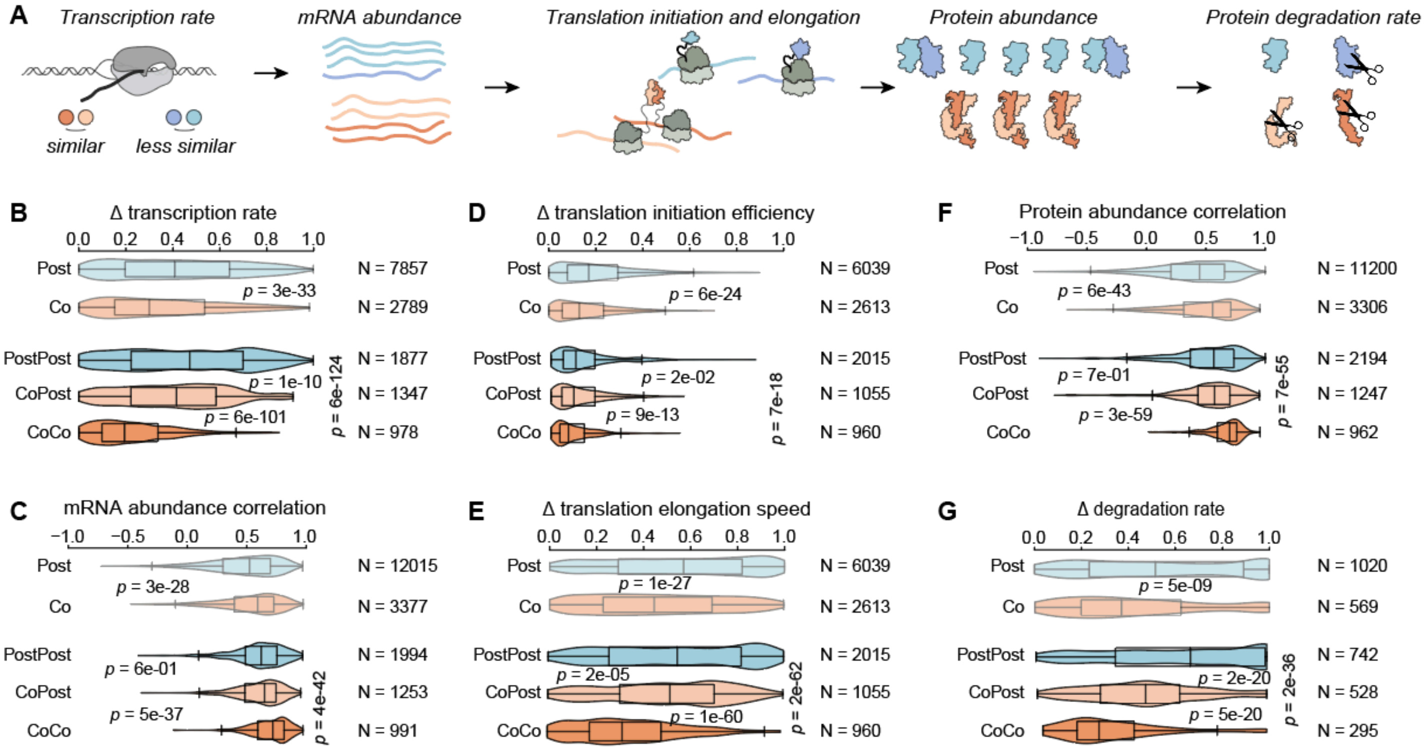


**Fig. S15. Violin plots Replicating Fig. 2, but after incorporating the directional component of co-translational assembly for human heteromeric pairs with experimental/model structures.**
(**A**) A schematic representation of various stages of the central dogma, same as Fig. 2A. (**B**) Violin plots comparing the gene transcription rate deviations of co- (orange) and post-translational (cyan) heteromeric co-complex pairs, and that of random pairs (gray). Plot features follow Fig 1B. The top two transparent violins are the same as Fig. 2B, without invoking structural information. The bottom three opaque violins include only those heteromeric pairs that appear in the same structure, and thus the directionality of assembly could be invoked. (**C**) Same as (B), but for Spearman correlations of mRNA abundances. (**D**) Same as (B), but for mRNA translational initiation efficiency deviations. (**E**) Same as (B), but for mRNA translational elongation speed deviations. (**F**) Same as (B), but for Spearman correlations of protein abundances. (**G**) Same as (B), but for protein degradation rate deviations.

Interestingly, the synchronization of the rates of core cellular processes of copost pairs appear to be intermediate to coco and post-post, but often more similar to the latter. This is consistent with the expectation that synchronizations of transcription, translation, and protein degradation are unlikely when one subunit is pre-translated and also stable in isolation but the other one is co-translationally dependent on it.

#### Note 5. Characterizing co-translational assembly of Acc1 homomer

We examined co-translational assembly of Acc1 homodimer by SeRP. To that end, we analyzed ribosomes co-purified with C-terminally tagged full-length Acc1. We noted increased ribosome footprint density onwards codon 250, which coincides with the synthesis and exposure of the homomeric interface (**Fig. 4B**). This result was consistent with our prediction that this homodimer assembles co-translationally.

However, the enrichment of ribosomes around codon 250 indicated high levels of interaction fluctuations. These fluctuations are in fact consistent with the intricate allosteric regulation of Acc1, leading to major conformational changes at the subunit interface (*4*). These results suggest a tantalizing possibility that allosteric regulation of the Acc1 interface may occur in a co-translational manner, which is also consistent with this enzyme's translation regulation (*5*).

#### Note 6. Interface asymmetry of *Ski2/3* and *Rrn6/7* affects co-translational assembly detection

We examined three predicted coco pairs by RIP-qPCR; those include Rpb1/2, Ski2/3, and Rrn6/7. For the Rpb1/2 pair, we found evidence that either subunit co-translationally engages with its full-length partner (**Fig. 4C**). However, for Ski2/3 we observed that translating Ski2 co-translationally engages with full-length Ski3, but not *vice versa* (**Fig. 4C**). Similarly for Rrn6/7, translating Rrn7 was observed to be co-translationally engaging with full-length Rrn6, but not *vice versa* (**Fig. S16A**).

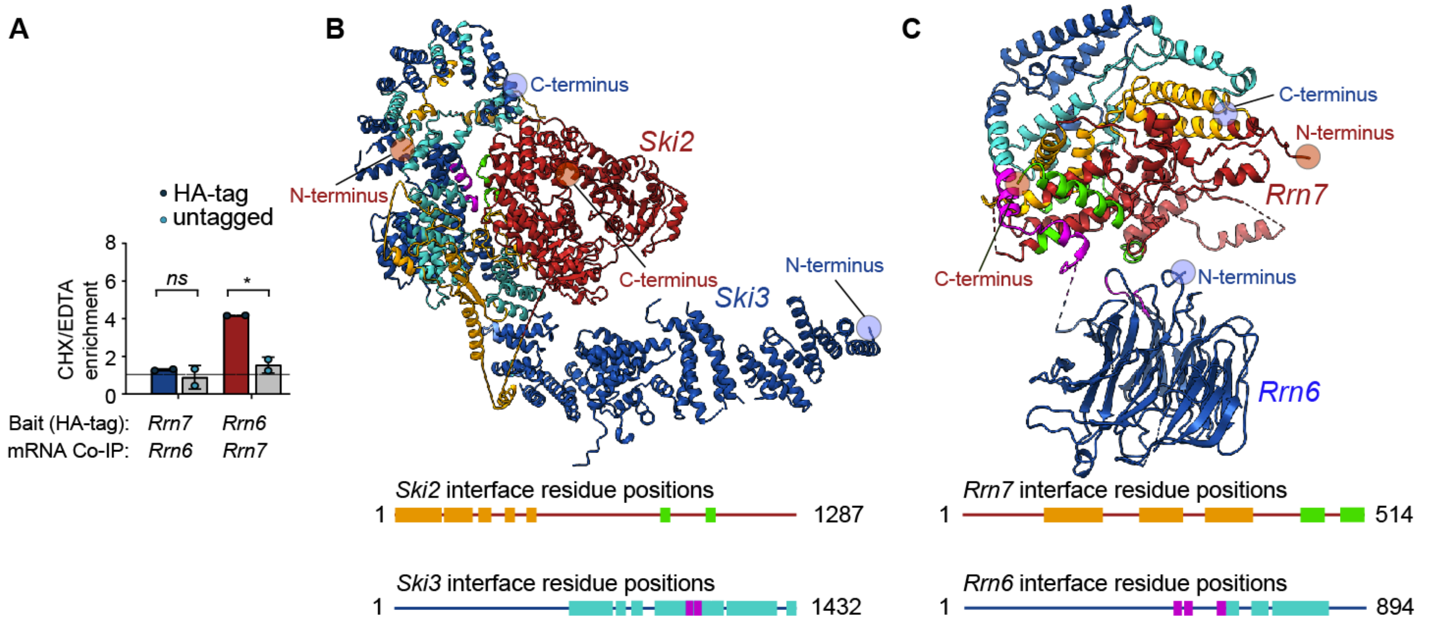


**Fig. S16**. **(A)** Quantifying co-translational assembly by RIP-qPCR for Rrn6/7. Barplot features are the same as **Fig. 4C**. **(B)** Cartoon view of the Ski2/3 dimer (PDB code: 4BUJ, ref.(*6*)). Ski2 is in red, Ski3 in blue; the interface residue positions are highlighted in the structure (top) and also represented as boxes (bottom). The N-terminus of Ski2 binds to the C-terminus of Ski3, and the respective interface residues are highlighted in orange and teal. The C-terminus of Ski2 also makes a few contacts (green residues) with Ski3 (magenta residues), which do not appear to be critical for the stability of the complex. **(C)** Same, for Rrn6/7 dimer (PDB code: 5O7X, ref.(*7*)).

These discrepancies may originate in the interface of one subunit being much closer to the C-terminus than the other subunit, allowing the former being more likely to be released from the ribosome first and be pulled down. As depicted in **Fig. S16B**, the N-terminus of Ski2 binds to the C-terminus of Ski3. In a co-translational scenario, once the intermolecular contacts crucial to stabilize the dimer (interface residues highlighted in orange for Ski2, teal for Ski3) are formed, Ski2 has nearly three-quarters of its length yet to be translated, while Ski3 translation is about to finish. This structural asymmetry is consistent with RIP-qPCR detecting translating Ski2 co-translationally engaging with full-length Ski3, but not *vice versa*.

Similar structural asymmetry exists in the Rrn6/7 pair, where the bulk of the interface comprises the N-terminus of Rrn7 and the C-terminus of Rrn6 (interface residues highlighted in orange and teal respectively, **Fig. S16C**). The C-terminus of Rrn7 indeed also contacts Rrn6 (interface residues highlighted in green and magenta). But, upon a critical visual inspection, these contacts do not appear to be critical for the stability of the dimer in a co-translational scenario. Hence, the structural asymmetry of Rrn6/7 also is consistent with RIP-qPCR detecting translating Rrn7 co-translationally engaging with full-length Rrn6, but not *vice versa*.

#### Note 7. Visualizing mRNA co-localization by smFISH

Detecting co-localization of mRNAs by smFISH (*Supplementary Methods 8*) involves two sets of fluorescent probes specifically binding to the two target mRNA populations in the cell. The efficiency of co-localization detection depends on (i) the probe-mRNA binding specificity, (ii) what fraction of the available mRNA molecules were bound to the probes, and (iii) the difference in their expression levels. What is the upper limit of co-localization detection if these factors were mitigated? To address this question, we examined two idealized scenarios: co-localization of the 5′ and 3′ halves of Rpb1 and Rpo31 mRNAs (positive controls). Both are constitutively expressed long mRNA molecules that are easy to target by probe-sets. As negative controls, we also examined the co-localization of each of them with unrelated mRNAs (Nuf2 and Ret1). As plotted in **Fig. S17**, for Rpb1, nearly 72% of all detection spots (red + green spots) were co-localized (yellow), and for Rpo31, that number was only about 47%. Almost no co-localization signal was detected in the negative controls.

These results indicate that even in the case of 100% biological co-localization, by smFISH method only about 50-70% can be detected. For two different mRNAs the biological co-localization is expected to be mitigated by the fraction of actively translated mRNAs and potential alternative assembly pathways not involving co-translational assembly, both of which are unknown. Hence, smFISH methods are expected to detect relatively lower co-localization measures for predicted coco heteromeric pairs (**Fig. 4D**).

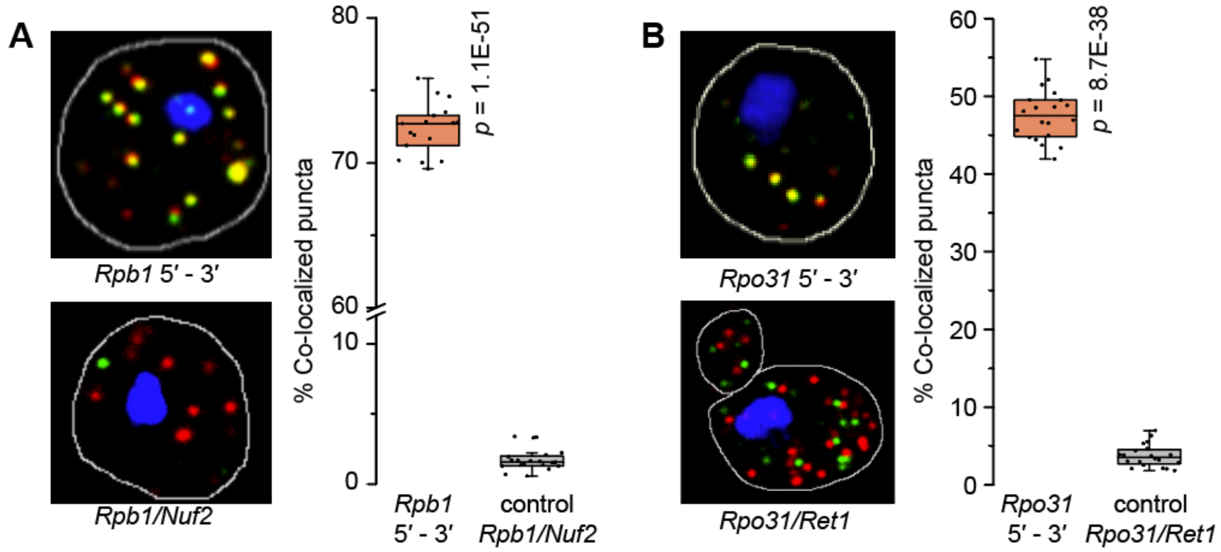


**Fig. S17**. RNA-smFISH experiments in yeast cells depict specific co-localization of the positive controls (5′ and 3′ halves of *Rpb1* and *Rpo31*). For the positive controls, the green and red spots correspond to the 5′ and 3′ halves of the same mRNA. For the negative controls, they represent two different mRNAs. For each pair, box plots represent quantified co-localization (%co-localized spots out of all red+green spots) over all cells in the field-of-view. As controls, we tracked the co-localization of each target mRNA with those of unrelated genes (*Nuf2* and *Ret1*). (**A**) Comparing co-localization of the 5′ and 3′ halves of *Rpb1* mRNA, as compared to the co-localization between *Rpb1* and *Nuf2*. (**B**) Comparing co-localization of the 5′ and 3′ halves of *Rpo31* mRNA, as compared to the co-localization between *Rpo31* and *Ret1*.

The list of predicted coco heteromeric pairs included in the smFISH experiments are enlisted in **Table S4**. These pairs correspond to complexes with unrelated functions, and therefore, any two proteins from two different complexes are not expected to show co-localization and could be used as controls.

**Table S4**. Enlisted are the positive control, predicted co-translational heteromeric pairs, and negative control pairs tested by smFISH experiments for mRNA colocalization.

| **smFISH pairs tested (predicted coco heteromers, co-localization expected)** | **control pairs (pairs of unrelated mRNAs, co-localization not expected)** |
|---|---|
| Positive Control: 5′ and 3′ halves of Rpb1 mRNA | Rpb1 vs Nuf2 |
| Positive Control: 5′ and 3′ halves of Rpo31 mRNA | Rpo31 vs Ret1 |
| Predicted coco pair: Rpb2 vs Rpb1 (RNA polymerase-II) | Rpb2 vs Ndc80, Rpo21 vs Nuf2 |
| Predicted coco pair: Rpo32 vs Ret1 (RNA polymerase-III) | Ret1 vs Gcd14, Rpo31 vs Yku70 |
| Predicted coco pair: Ski2 vs Ski3 (RNA helicase) | Ski3 vs Yku80, Ski2 vs Gcd10 |
| Predicted coco pair: Gcd10 vs Gcd14 (tRNA methyltransferase) | Gcd10 vs Yku80, Gcd14 vs Yku70 |
| Predicted coco pair: Yku70 vs Yku80 (kinetochore) | Yku70 vs Rpo31, Yku80 vs Ski3 |
| Predicted coco pair: Nuf2 vs Ndc80 | Nuf2 vs Rpb1, Ndc80 vs Rpb2 |

Four out of six pairs displayed statistically significant signatures of mRNA co-localization (**Fig. 4D**).

The results for the remaining two pairs are shown in **Fig. S18**.

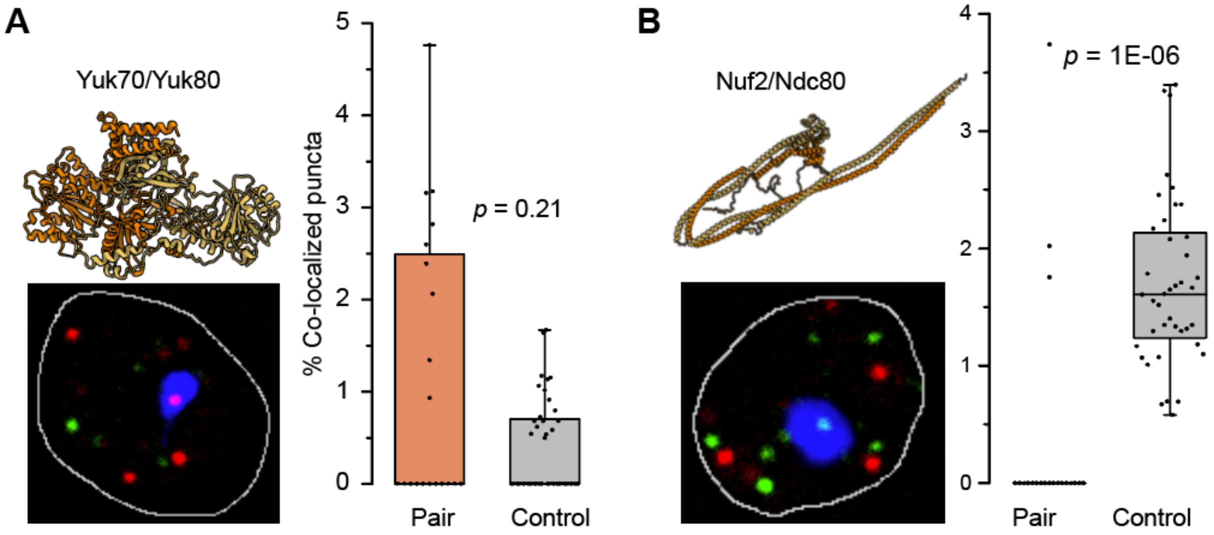


**Fig. S18**. RNA-smFISH experiments in yeast cells attempting to find signatures of specific co-localization of mRNAs of two predicted coco heteromeric pairs, (**A**) Yuk70/80, and (**B**) Nuf2/Ndc80. No significant co-localization signals were obtained for these two pairs. Control experiments tested their co-localization with unrelated mRNAs (Yku70 vs Rpo31, Yku80 vs Ski3) and (Nuf2 vs Rpb1, Ndc80 vs Rpb2). Figure specifications follow Fig. S19.

#### Note 8. Protein abundance drop upon genetic knockout of co-translational partner

Genetic perturbation experiments depicted in **Fig. 4E** show that compared to post, co-translational heteromeric proteins tend to be relatively more unstable in isolation and tend to form aggregated foci visible in fluorescent microscopy.

To obtain a more generalized picture, we analyzed a recently published dataset in which the abundance drop of thousands of yeast proteins in the soluble fraction were quantified by mass spectrometry upon single gene knockouts (*8*). If a co-translational protein is partner-stabilized, genetic knockout of the partner is expected to cause a drop in its abundance. To test that we first analyzed seven predicted copost pairs (**Table S5**).

**Table S5**. Predicted copost pairs. The 3D structures of these pairs were originally predicted by AlphaFold (see ref.(*9*)). A→B assembly means A assembles on B as it is being translated. B is pre-translated and presumably stable in isolation.

| **Gene1** | **assembly direction** | **Gene2** | **ModelArchive code,** ref.(*9*) |
|---|---|---|---|
| SNF1 | → | SNF4 | ma-bak-cepc-0074 |
| CKB1 | → | CKA1 | ma-bak-cepc-0144 |
| YMR31 | → | KGD2 | ma-bak-cepc-0168 |
| EMP47 | → | SSP120 | ma-bak-cepc-0599 |
| BUG1 | → | GRH1 | ma-bak-cepc-0891 |
| STO1 | → | CBC2 | ma-bak-cepc-0991 |
| MDY2 | → | GET4 | ma-bak-cepc-1016 |

For these seven A→B unidirectional pairs, we computed the $\log_2$ fold-change as compared to knockout/unperturbed conditions, of A upon B knockout, and of B upon A knockout. We found significantly larger abundance drops in the former, compared to the latter (**Fig. S19A**).

We extended this analysis to all predicted co- and post-translational pairs in yeast, for which the abundance of at least one protein was available, upon knockout of its partner. As plotted in **Fig. S19B**, predicted co-translationally assembling proteins suffered significantly larger abundance drops than subunits than those predicted to be stable in isolation.

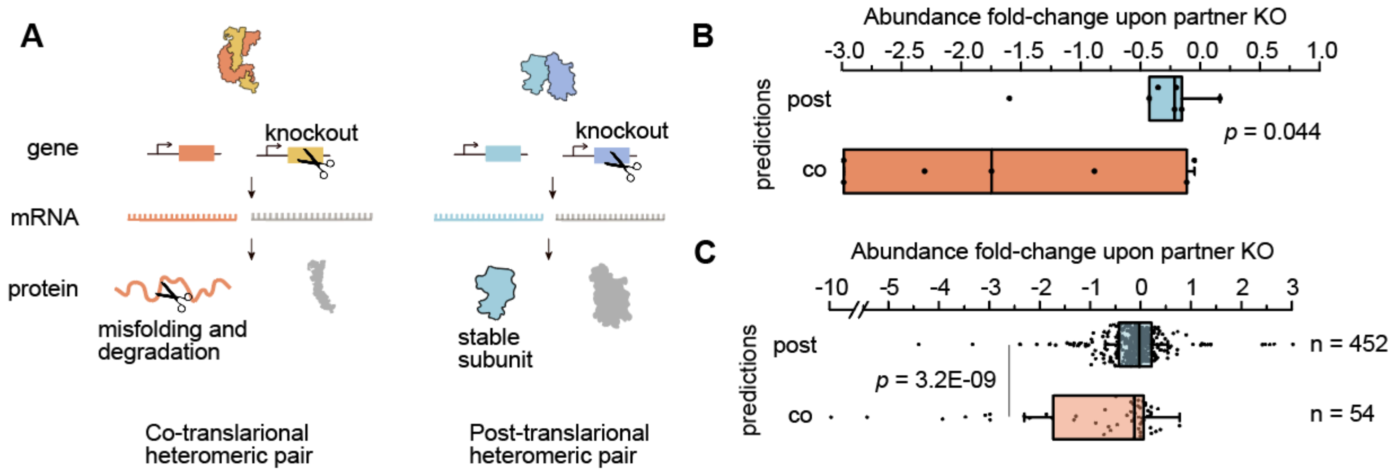


**Fig. S19**. (**A**) A schematic representation of the effects of partner knockout on co- and post-translational pairs. Because inter-protein contacts are crucial to the stability of co-translational proteins, partner gene knockout is expected to cause misfolding and degradation, and thus a drop in the abundance as compared to unperturbed conditions. This abundance drop is not expected for a post-translational pair, in which each subunit is expected to be stable in isolation. (**B**) Boxplots depicting the distribution of protein $\log_2$ abundance fold-change upon partner knockout in co- (orange) and post-translational (cyan) proteins predicted in yeast. This plot includes seven unidirectional pairs enlisted in Table S2, for which fold-changes were available for knockout of either protein. (**C**) Extending the analysis to 506 predicted yeast pairs, for which fold-changes were available for knockout of at least one protein.
